# The Omniroute maze: a novel rodent navigation apparatus that integrates dynamically configurable routes, sensory cues and automated reward delivery

**DOI:** 10.1101/2025.09.01.672969

**Authors:** Adam W. Lester, Afsoon G. Mombeini, Manu S. Madhav

## Abstract

We developed the Omniroute maze, a novel apparatus that enables near-fully automated, high-throughput behavioral experiments in freely moving rodents, combining the experimental control of virtual reality (VR) platforms with the ethological validity of real-world navigation. The maze features movable wall segments that can be programmatically configured to create unique routes within a 90 × 90 cm platform. Four projectors arrayed around the maze perimeter can display distinct visual cues on both sides of any subset of raised gate wall panels and the maze floor as well as play directional auditory cues. A motorized gantry delivers food-based rewards anywhere in the maze. Real-time 3D tracking of the rat and the reward gantry enables closed-loop control of maze topology, cue configuration, and reward delivery. All subsystems are controlled using the Robot Operating System (ROS) framework via a custom Python-based user interface. The Omniroute can reproduce classic behavioral mazes and a variety of novel configurations to test hypotheses on the interactions between routes, cues, and behavior. Automated configuration, tracking, and reward delivery enable high-throughput experiments on complex navigation behaviors without the potential biases introduced by direct experimenter intervention. Designed from the ground up for robust operation, the Omniroute system utilizes affordable hardware and software to facilitate easy fabrication and assembly as well as replicability by other researchers.

**Significance Statement:** The Omniroute system enables high-throughput, automated rodent 2D navigation experiments with real-time control over routes, cues, and reward delivery, comparable to that afforded by modern virtual reality-based systems. In addition, however, the Omniroute supports uninterrupted movement-related behavior, which is critical for assessing the ongoing influence of self-motion cues, while still minimizing confounds from experimenter handling. Naturalistic food-based reinforcement at any location allows for closed-loop behavioral conditioning under diverse task structures. The Omniroute system is one of very few platforms that support near-complete control of available paths, dynamic visual and auditory cues, and naturalistic food-based reinforcement with minimal direct experimenter intervention.

## Introduction

Route-based navigation tasks remain foundational in behavioral neuroscience, offering a window into spatial memory, decision-making, and cognitive flexibility in rodents. From canonical physical mazes like the radial arm maze (Olton and Samuelson, 1976) to more recent rodent virtual reality systems (Dombeck et al., 2007; Ravassard et al., 2013; Thurley and Ayaz, 2017), maze-based navigation tasks have enabled detailed investigations of hippocampal function (O’Keefe and Dostrovsky, 1971; Muller et al., 1987), place learning (Packard and McGaugh, 1996), and neural representations of space (O’Keefe and Nadel, 1978; Wilson and McNaughton, 1993). Despite the widespread adoption of mazes in rodent research, however, most remain limited in their application, and often require direct experimenter involvement to alter route access, modify stimuli, or deliver reinforcement. These manual interventions limit the throughput and repeatability of experiments. They can also introduce unwanted stress, olfactory cues, experimenter bias, and interruptions to ongoing behavioral and cognitive operations (Lester et al., 2020).

Open-source ecosystems have lowered barriers for investigators to build and incorporate custom semi-automated behavioral systems into their experimental arsenal, sidestepping these limitations to a degree (Lopes et al., 2015; White et al., 2019; Jankowski et al., 2023). Few of these open options, however, support the types of automated operation that are available with more expensive proprietary alternatives (Holleman et al., 2019; Sawatani et al., 2022; Jankowski et al., 2023; Porter et al., 2025). Furthermore, both open and proprietary systems are largely limited in their configurability. Advances in VR-based animal behavioral systems, in contrast, have enabled even more complete, precise, programmable control of environmental features and reinforcement. Importantly, however, most VR approaches inevitably decouple animal locomotion from proprioceptive and vestibular inputs (Chen et al., 2013; Ravassard et al., 2013; Aghajan et al., 2015), as they typically require animals to be head-fixed. These considerations have motivated more hybrid approaches that preserve real-world movement while allowing dynamic sensory control in physical environments (Jacobson et al., 2014; Kaupert et al., 2017; Stowers et al., 2017; Lester et al., 2020; Madhav et al., 2022), particularly for studies assessing the physiological mechanisms that subserve navigation. To our knowledge, however, none of the existing open, proprietary, VR or hybrid systems simultaneously support continuous 2D locomotion, real-time control of routes, multimodal cues, and closed-loop, spatially targeted reinforcement.

The Omniroute system was designed to support the flexibility and control of VR while preserving naturalistic real-world behavior. It is a predominantly open-source, modular platform that integrates unconstrained real-world navigation with rapid dynamic control over route topology, sensory stimuli, and reward delivery. Unlike conventional mazes or fixed hybrid systems, the Omniroute supports rapid reconfiguration of the environment in response to ongoing behavior, allowing for experiments that demand flexible reconfiguration of task constraints and real-time feedback.

The Omniroute system consists of a 90 × 90 cm maze with 60 independently movable motorized gates that allow for on-the-fly route reconfiguration. Four 4K short-throw projectors deliver visual cues onto maze wall and floor surfaces as well as directional auditory cues. A motion capture system tracks the rat and an XY gantry, which dispenses liquid reward at arbitrary maze locations. All components are coordinated through the Robot Operating System (ROS), enabling real-time rat position estimation and closed-loop control over gate position, cue configuration, and reward timing. This configuration supports a wide range of navigational and perceptual tasks. Task logic is implemented in software and remains decoupled from the physical layout, allowing the same hardware to instantiate diverse behavioral motifs from plus mazes to multi-goal foraging arenas. Modular design using accessible modern fabrication techniques, stock materials, standard fasteners, and open PCB designs supports its adaptation and customization by other investigators with the requisite technical expertise and resources. All design files, firmware, and control software are openly available and designed for accessibility, reproducibility, and integration with standard neuroscience toolchains.

In the following sections, we describe the design, integration, and validation of the Omniroute system. We provide detailed characterizations of each subsystem and demonstrate the system’s suitability for scalable, high-throughput, and ethologically valid behavioral neuroscience applications.

## Materials and Methods

### Subsystem Coordination and Integration Overview

The following section provides a high-level overview of the major subsystems and their integration into a unified control architecture, with further details presented in subsequent sections and summarized in Figure 2.

#### Constituent Subsystems

The Omniroute integrates four hardware subsystems (Figs. 1 and 2).

1. **Gate Control**: Programmable gates provide reconfigurable routes using 60 motorized gate modules, each with a custom motor driver PCB (Figs. 3, 4).
2. **Stimulus Delivery:** Visual and auditory stimuli are delivered by four projectors that render floor and gate wall panel images and use integrated speakers for audio playback (Fig. 5).
3. **Reward:** A gantry enables XY positioning and liquid food reward dispensing utilizing a servo-actuated cannula and a peristaltic pump (Fig. 6).
4. **Tracking:** A multi-camera system that streams real-time 3D poses for the rat, the gantry, and maze perimeter reference markers (Fig. 7).

**Figure 1.**
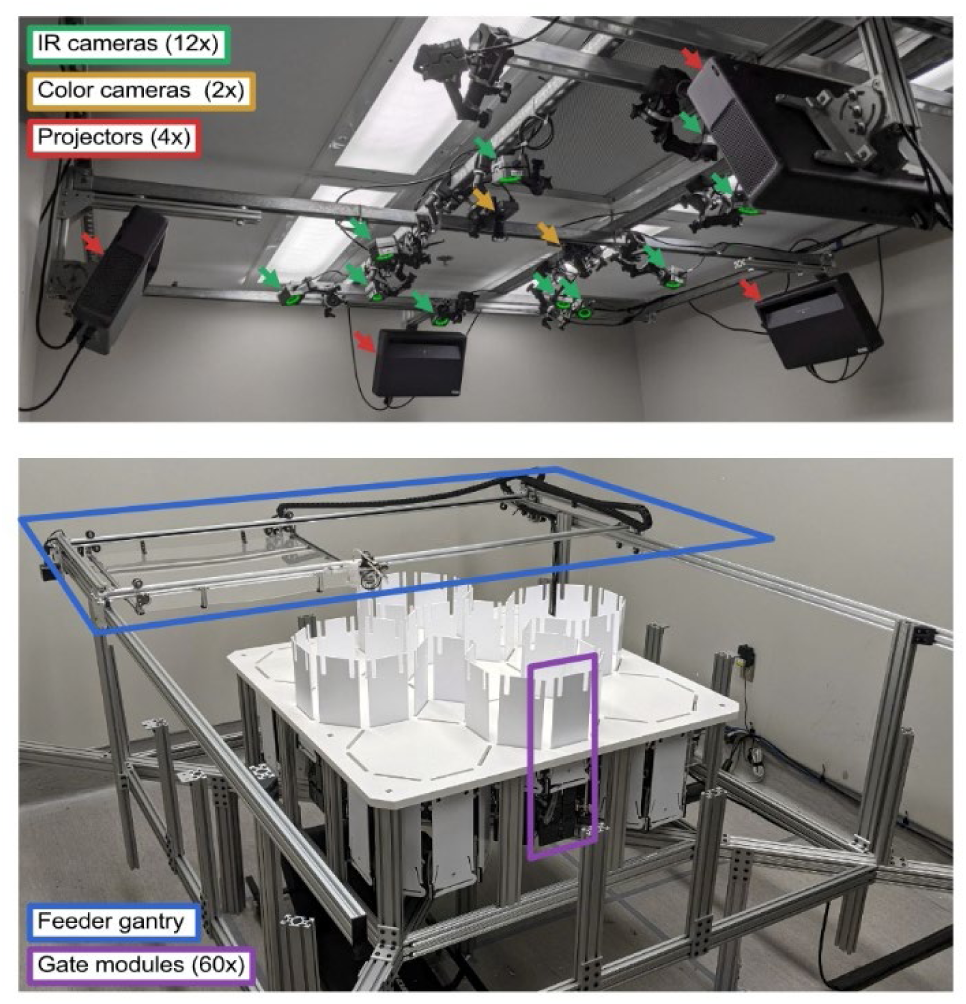
Omniroute system overview. (top) Stimulus delivery and position tracking infrastructure. This includes 12 infrared cameras (green), which provide real-time positional tracking, and 2 color cameras (yellow). Also shown are 4 laser projectors with integrated speakers (red) used to deliver visual and auditory cues. These components are mounted to a modular ceiling frame to provide stable, comprehensive coverage of the maze. (bottom) Maze with gate modules (purple). The 3×3 maze grid comprises nine octagonal chambers with 60 gates, enabling dynamic reconfiguration of routes. Also shown is the gantry system (blue), which delivers food reward at any location via an actuated cannula.

**Figure 2.**
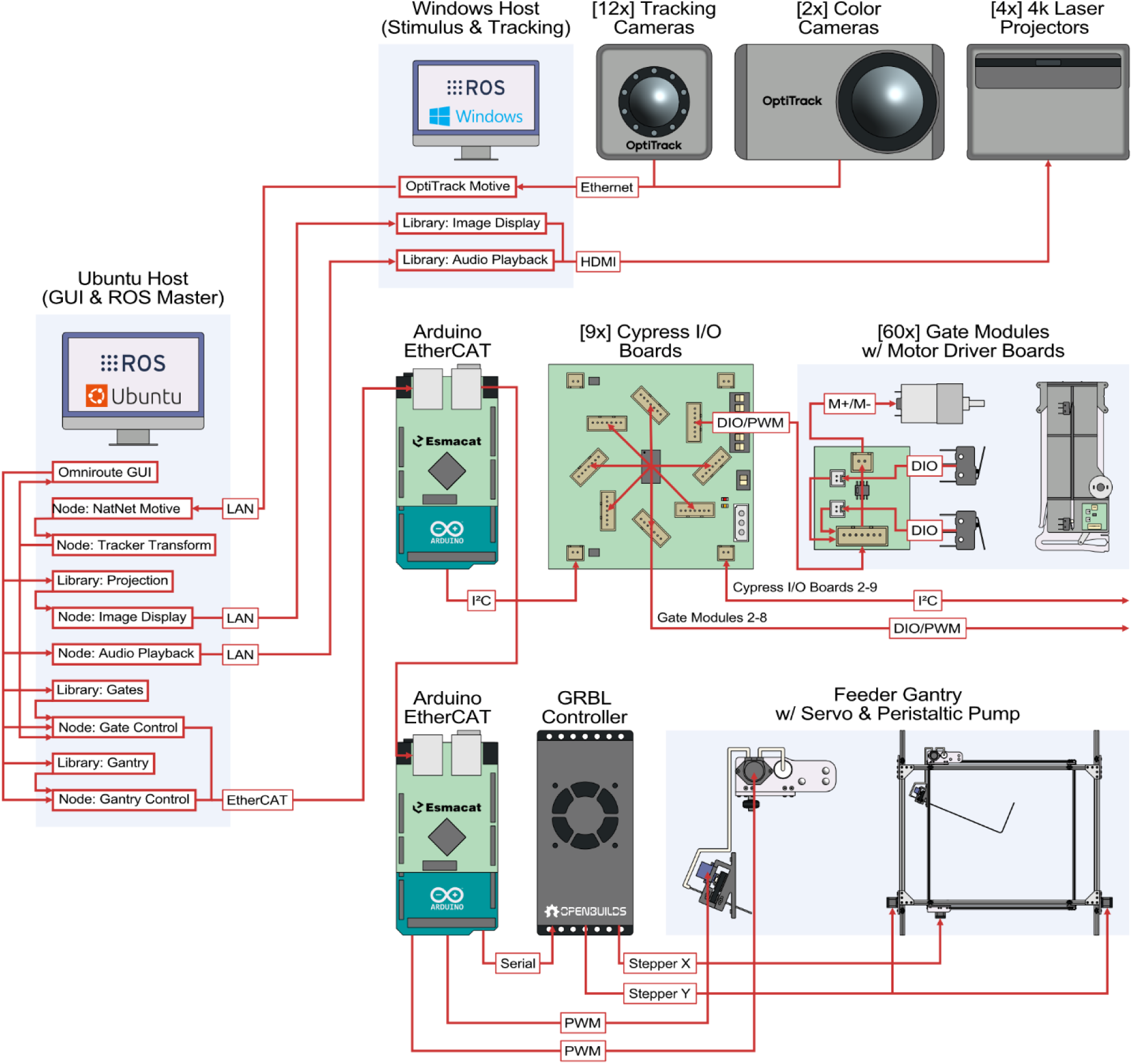
Schematic of the Omniroute system. The full system comprises multiple hardware subsystems coordinated via the ROS framework using nodes on two host computers. An Ubuntu host acts as the ROS master and manages the GUI and high-level commands for gates and gantry. The Windows-based host handles tracking and projector-based stimulus display. Subsystems include 60 gate modules actuated via custom motor driver boards and controlled through a network of Cypress I/O expanders and EtherCAT-linked Arduinos; a gantry system operated via GRBL controller and servo-driven pump; four projectors and integrated speakers managed via display and audio nodes; and a 14-camera tracking system (12 IR and 2 color cameras) that streams 3D position data and video for real-time closed-loop control.

#### Control Hosts and ROS Framework

Control of the subsystems is provided by the Robot Operating System (ROS) (Quigley et al., 2009), which is a software framework originally developed for robotics that allows multiple independent programs (nodes) to communicate to each other within the same computer and across computers on a network, by writing data (messages) to shared channels (topics). Our control logic is divided across ROS nodes running on two computers. A Unix-based computer runs the ROS master node and child nodes that handle central behavioral control logic and control multiple subsystems. This computer also hosts the graphical user interface (GUI) for system configuration, live control, and state visualization. A Windows-based computer runs the tracking system as well as projection display and audio playback nodes.

#### Gate and Reward Control Pipeline

For the gate control and reward subsystems, low-level hardware actuation is managed via two separate Arduino Mega 2560 microcontrollers (Arduino S. r. l., Monza, Italy) using Esmacat Arduino Shields (EASE; Harmonic Bionics Inc., TX, USA). These shields support daisy-chained Power-over-Ethernet (PoE) for power and the EtherCAT protocol for data, minimizing cabling complexity while supporting high-speed data transfer. An open-source library (Harmonic Bionics Inc., TX, USA) allows communication with the EASE boards through ROS. Gate state feedback and gantry actuation status are published to ROS topics to support closed-loop synchronization with other subsystems and ensure robust real-time operation during task execution.

#### Stimulus Delivery and Tracking Pipeline

Sensory stimulus delivery is handled on the Windows computer, with ROS nodes controlling image projection and audio playback from four short-throw projectors. Visual content is mapped to specific floor and gate wall panels using precomputed calibration data and can be updated dynamically based on trial context or the animal’s behavioral state. A multi-camera system (OptiTrack, NaturalPoint, Inc., Corvallis, OR, USA) provides real-time 3D tracking of both the rat and gantry. Custom ROS nodes transform this data into a shared maze-centered coordinate frame, enabling behaviorally contingent control of stimuli, gates, and reward targeting.

### Gate Control Subsystem

The Omniroute incorporates 60 actuatable gate modules that dynamically reconfigure route topologies during behavioral tasks. Each gate can be independently raised or lowered under software control at any time, enabling the creation of diverse maze configurations within a 3 × 3 octagonal grid (Fig. 3). The gates are distributed uniformly across the maze’s surface and the octagonal geometry supports both orthogonal and diagonal path trajectories between maze chambers. This component of the system has been described in great detail previously, along with the custom motor driver and I/O expander PCBs (Lester et al., 2024). Below is a brief overview of this component of the system, including details related to the integration of the gate modules into the larger Omniroute system.

**Figure 3.**
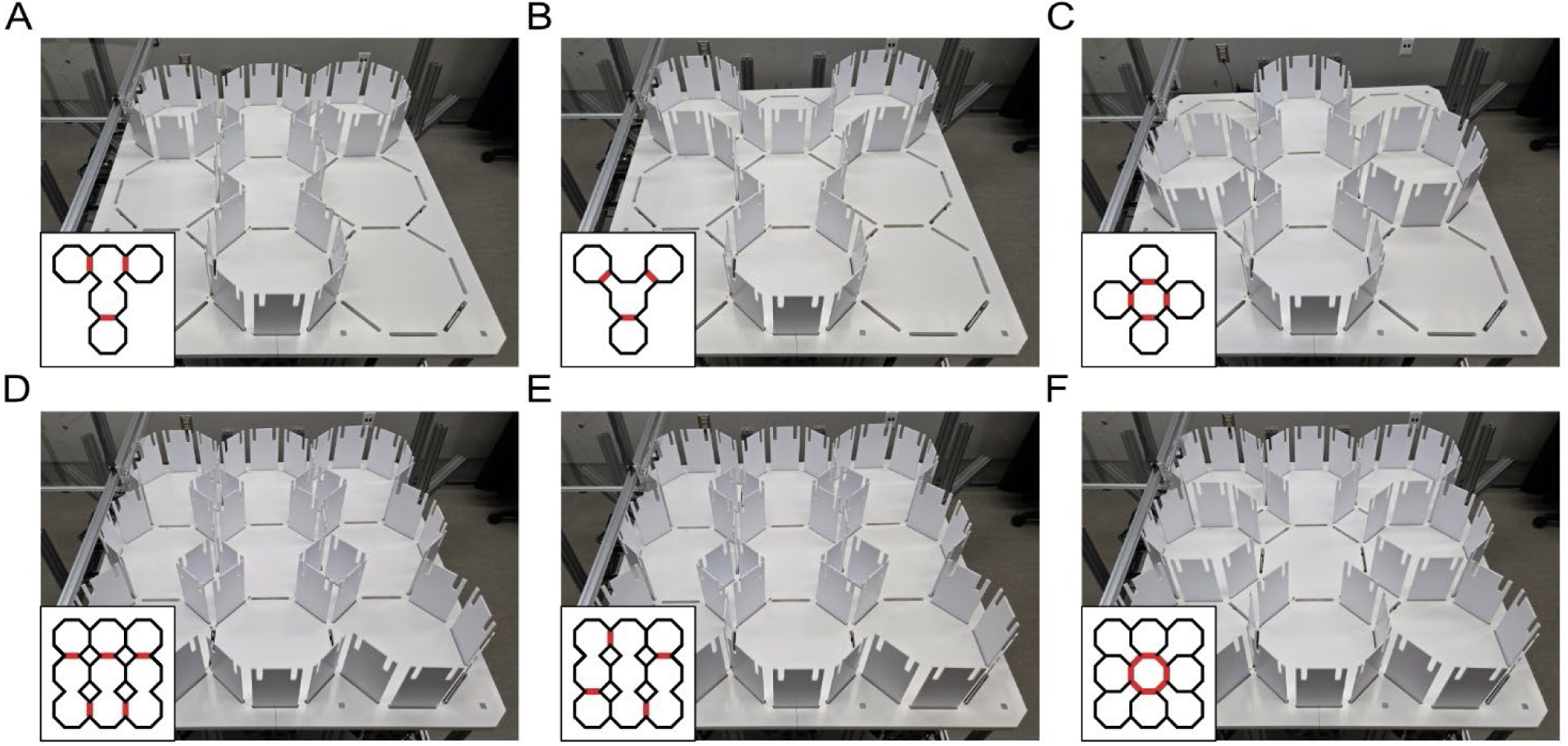
Example route configurations. Each panel shows a distinct gate configuration that mimics mazes commonly used in rodent behavioral experiments, achieved via programmatic control of the 60 gate modules. Insets depict schematic top-down representations of the static gate configurations (black) as well as potential dynamically controlled gates that may be used in these paradigms (red). The examples include **(A)** T-maze, **(B)** Y-maze, **(C)** plus maze, **(D)** W-maze, **(E)** hairpin maze, and **(F)** 8-arm radial maze. This demonstrates the system’s versatility to replicate canonical behavioral apparatuses and highlights how dynamically configurable gates can be used to, for example, pause the animal within the start zone, goal zone, or any other location.

#### Mechanical Architecture

Gates consist of a 100 mm wide wall panel driven vertically via a 12 V, 50 rpm, geared DC motor (FIT0492-A, DFRobot, Shanghai, China) coupled to a waterjet-cut pin-in-slot aluminum linkage (Fig. 4A and B). The assembly is built from laser-cut acetal panels with snap-fit joints, allowing for rapid assembly using a minimum of standard hardware and fasteners. Gate travel is ∼180 mm with limit switches detecting both upper and lower positions. The current configuration supports a density of up to eight gates per maze chamber, allowing tight spatial control of inter-chamber access.

**Figure 4.**
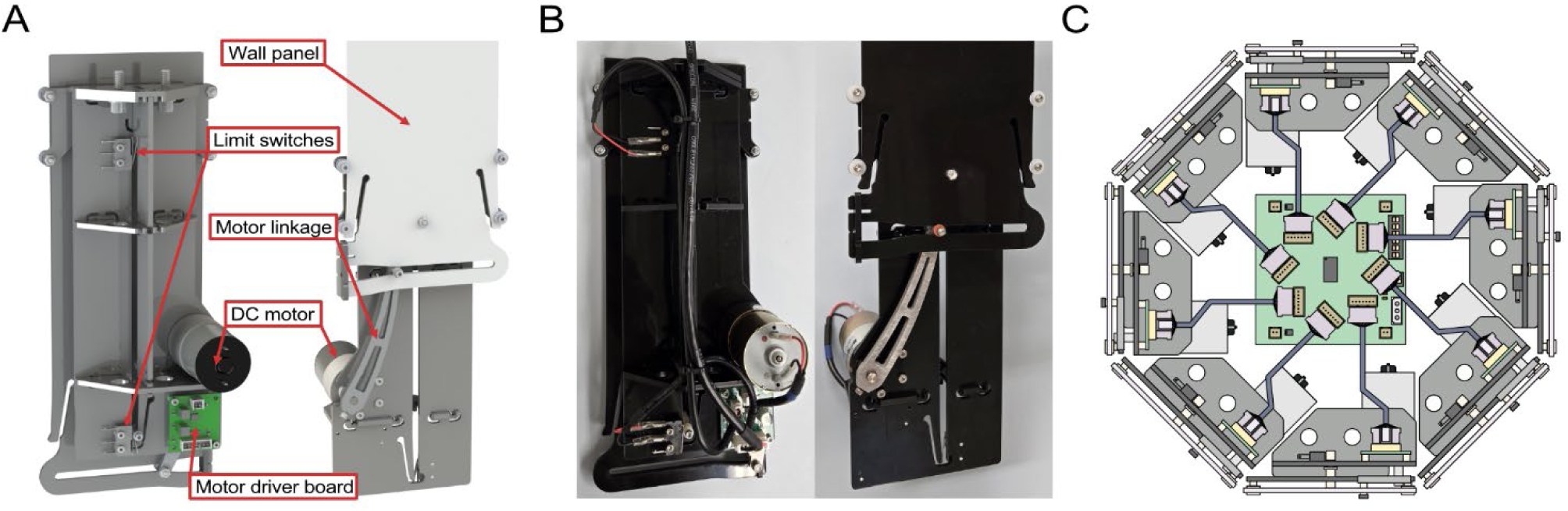
Gate module hardware and wiring topology. **(A)** CAD renderings of the front and back of a gate module, illustrating the key components. The gate wall panel undergoes vertical translation with a total travel of 180 mm. Limit switches detect the upper and lower travel endpoints. A pin-in-slot linkage converts rotary motion from a 12 V, 50 rpm geared DC motor into linear displacement of the gate wall panel. The custom motor driver board provides PWM-based motor control via an H-bridge and returns limit switch signals to the Cypress I/O board. **(B)** Photographs of a fully assembled gate module, including complete wiring. **(C)** Schematic of the gate module wiring. A total of up to eight gates interface with a single Cypress I/O expander board via PWM and digital I/O lines, carried by a 7-pin cable, supporting distributed control over all 60 gates through a buffered I²C network. **(A)** and **(B)** were adapted from (Lester et al., 2024).

#### Maze platform and frame

The maze platform consists of 19.25-mm-thick, CNC-machined foam PVC panel mounted to a support frame constructed from 80/20 T-slot aluminum profiles (Fig. 1). The base panel includes slots for the 60 gate modules and heat-set threaded inserts for fastening the platform to the frame and mounting the modules from below, ensuring no fastening hardware is visible on the maze surface. The frame is sized for the current 3 × 3 octagonal chamber layout but is designed to enable extension to a 5 × 5 configuration (160 gates) in the future.

#### Actuation and Control Hardware

Gate movement is powered by dedicated custom motor driver boards mounted on each module. These PCBs contain an H-bridge circuit (DRV8870DDAR; Texas Instruments, Dallas, TX, USA) and transistor-based limit switch cutoffs to ensure rapid and reliable termination at the endpoints of travel.

Up to eight gates can be grouped and connected to a single custom Cypress-based I/O expander board (CY8C9540A, Infineon Technologies AG, Neubiberg, Germany), which handles gate control via I²C. Each gate module communicates with its respective Cypress board via a 7-pin connector carrying PWM control and limit switch state from the motor driver board (Fig. 4C). These Cypress boards provide enough independent PWM and digital I/O channels to manage all gates in the group and support address selection for up to 64 independent boards. While the current iteration of the Omniroute system utilizes only 9 Cypress boards and 60 gates, this design enables theoretical scaling to 512 gates using 64 Cypress boards and a single microcontroller.

Cypress boards communicate over a buffered I²C bus to a central Arduino Mega 2560, which acts as the main gate controller. The system uses single- or double-buffered configurations via P82B715 buffer chips (NXP Semiconductors, Eindhoven, Netherlands) to extend bus length beyond standard I²C constraints. This enables reliable communication across the spatially distributed layout of the 90 × 90 cm platform. All boards are powered by a 12 V (motor) and 5 V (logic) bus, with centralized supply and distribution via a modular power rail.

#### ROS Integration

The gate system is fully integrated into the Omniroute control architecture (Fig. 2). A dedicated gate control ROS node handles route configuration and emits control signals to the gate controller node on the Arduino. These commands are relayed via an EASE Shield to the Arduino, which parses movement schedules and actuates gates via I²C through the nine Cypress boards. Gate position state, based on limit switch feedback from each Cypress board, is published back to ROS and used to verify actuation success, support conditional routing logic, and synchronize movement with other subsystems (e.g., stimulus delivery and reward).

Gates can be actuated in closed-loop fashion based on live animal position, or pre-defined route layouts. Control messages support both individual gate operations and batched commands for system-wide route reconfiguration.

In combination with the high-speed tracking system, the gate control architecture allows for complex real-time interaction paradigms.

### Stimulus Delivery Subsystem

#### Projectors

Four short-throw Mi 4K Laser Projectors (XMJGTYDS01FM; Xiaomi Corp., Beijing, China) are positioned around the maze perimeter to provide full 360° coverage (Fig. 5A). These projectors support motorized focus and 8-point keystone correction and feature short-throw optics capable of covering the complete maze platform and gate wall panel surface. Each projector also includes four integrated speakers (2 × full-range + 2 × tweeters; 30 W total output).

**Figure 5.**
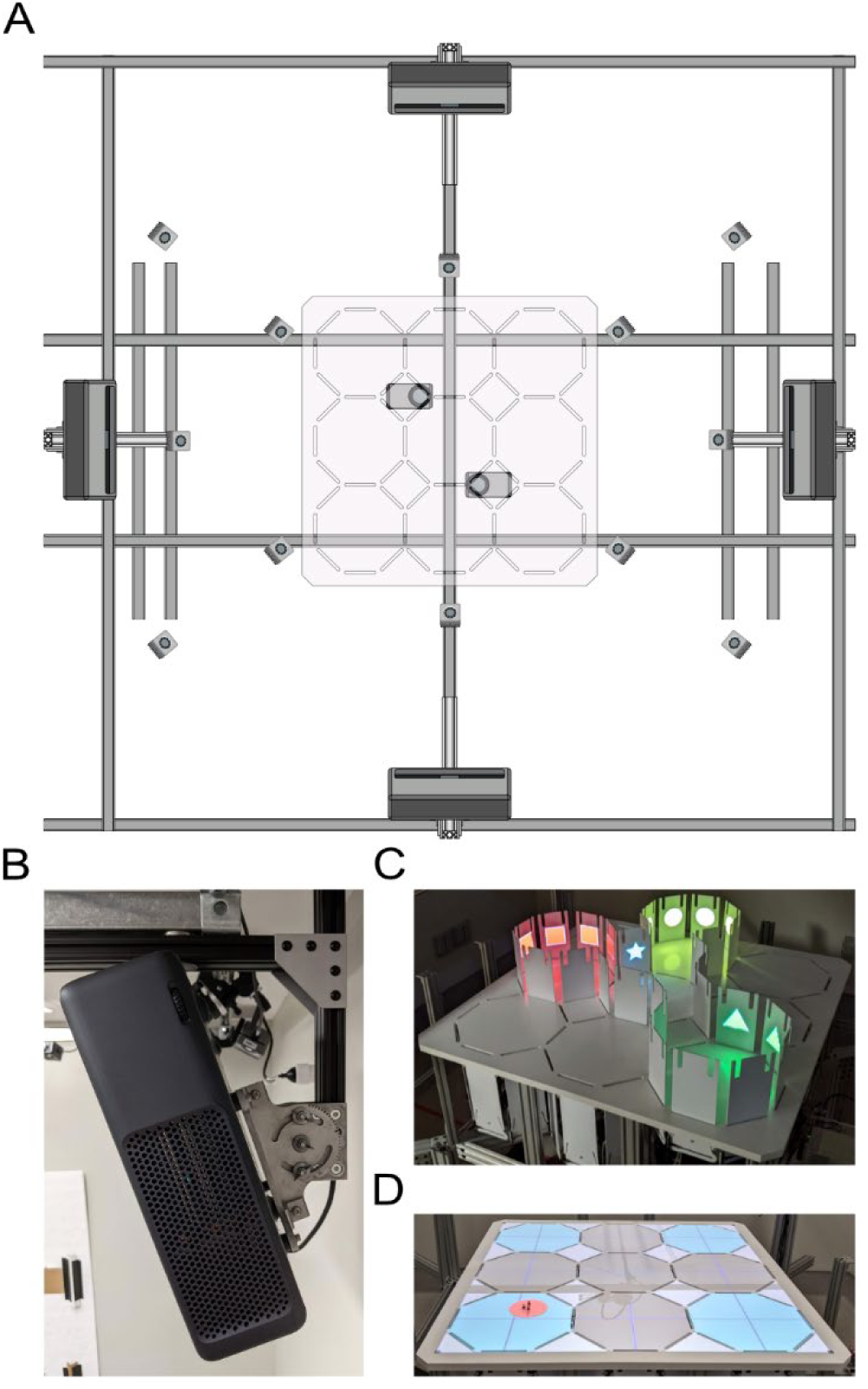
Visual and auditory stimulus delivery. **(A)** Bottom-up schematic showing the spatial arrangement of the four projectors (centered on each maze edge). Also shown, are the 12 IR cameras (angled inward from corners and sides), and the two color cameras positioned near the center axis of the maze. All components are mounted into a strut-channel ceiling frame, providing full coverage of the maze floor and gate wall panel surfaces. **(B)** Close-up of a projector mounted to a custom yaw-adjustable aluminum bracket. **(C)** Example projection of gate wall panel stimuli, with distinct images rendered on the inner faces of raised gate panels. **(D)** Example of projected floor stimuli, including a dynamic mask aligned with the rat’s position to avoid retinal exposure. This mask, shown in red for visibility, is black and 15 cm in diameter during actual experiments.

Projectors are mounted overhead using custom waterjet-cut mounts fabricated from 6.35 mm (1/4”) thick aluminum (Fig. 5B). These mounts provide mechanical stability while allowing fine-grained yaw adjustment over a ±45° range. The four projectors are arrayed around the maze at 90° intervals, with each mounted to a modular framing system composed of 80/20 extruded aluminum profiles anchored into Unistrut-style channels in the ceiling of the room housing the system. This arrangement enables consistent and complete angular coverage as well as unobstructed line-of-sight from each projector to its assigned projection zones (maze floor or gate wall panels).

#### Projection Calibration

Each projector requires a one-time initial calibration. Calibration is performed independently for each projector to ensure accurate image warping onto either the floor or vertically raised gate wall panel. Calibration is performed using a custom OpenGL-based ROS node that renders geometric test patterns to each projector and supports manual control point adjustment via keyboard input. The system uses normalized device coordinates (NDC) to represent both maze-space and display-space vertices, enabling projector-independent coordinate mapping.

During calibration, the user selects a calibration mode (floor or gate wall panels), after which they manually align four control points projected onto the maze with known physical reference points (i.e., the gate wall or maze floor corners). A homography matrix is then computed for each maze location by solving the least-squares minimization problem between source (image-space) and target (NDC-space) coordinates. This homography provides a transformation from source to target space for each projector and location. Homographies are saved per-projector to an XML file and reloaded at runtime for projection correction.

#### Visual cues

Visual stimuli can be projected onto either side of any of the 60 gates when they are in their raised configuration (Fig. 5C) or the maze floor (Fig. 5D). Each projector is controlled by a dedicated OpenGL context running in the ROS image display node on the Windows computer (Fig. 2). Projection image textures are stored as 8-bit RGBA PNG files, preloaded into memory, and mapped to projector coordinates at runtime using the precomputed homography matrices loaded from the calibration XML files, with separate matrices for each gate wall panel segment and floor region.

At runtime, ROS messages on the projection image topic specify which stimulus image to display at each gate wall panel and maze floor location. All rendering operations, including homography warping, texture loading, and shader-based drawing, are executed per-frame at the native refresh rate of the projectors (60 Hz), ensuring flicker-free display suitable for behavioral tasks.

In addition to static stimulus images, the system projects a 15 cm diameter mask around the rat’s head at its current position, removing the potential for retinal exposure to high-intensity light (Fig. 5D). The mask is computed in real time based on rat’s motion capture data and rendered using a circular shader warped into projector space using the appropriate homography for each projector.

#### Auditory Cues

Auditory stimuli are delivered through speakers integrated into each projector, using a ROS audio playback node (Fig. 2). Sound files are stored as WAV files and preloaded into memory at startup for low-latency playback. Each projector is a unique audio output device. By default, audio cues are played concurrently on all four projector speakers, but can be routed to a subset of projectors as needed. This architecture supports directional auditory cues and reliable synchronization of audio with visual cues or behavioral events in real-time.

### Reward Subsystem

#### Hardware

The reward delivery system is built around a modified gantry (ACRO, OpenBuilds, Zephyrhills, Florida, USA), designed to support reward delivery anywhere in the maze (Fig. 6A). The XY motion stage uses V-slot aluminum extrusions (20 × 40 mm profile) and Delrin V-wheels for linear motion, with three NEMA 17 stepper motors driving movement via GT2 timing belts. Mounted beneath the moving gantry carriage is a 610 mm × 610 mm, 1/6” (4.2 mm) thick acrylic ceiling panel that serves as a dynamic overhead enclosure to prevent the rat from climbing over raised gates (Fig. 6B). The panel is fixed to a 10 mm aluminum frame that also supports the food dispensing components.

**Figure 6.**
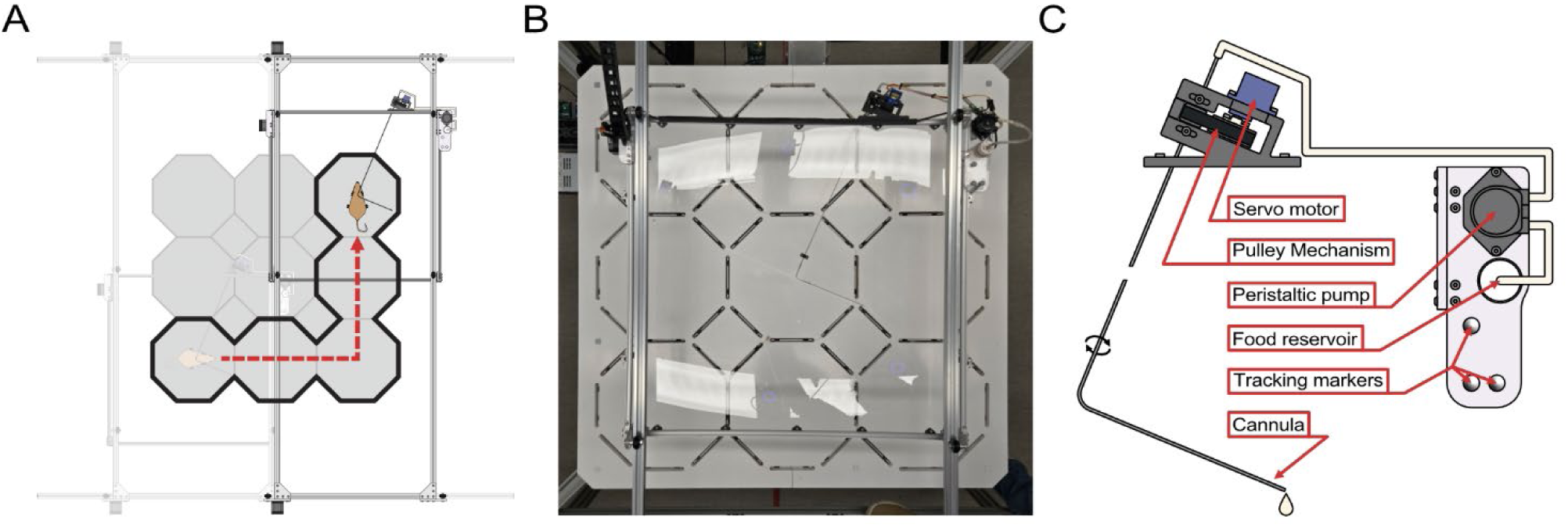
Automated reward delivery system. **(A)** Schematic showing the XY gantry system tracking the rat’s position in real-time to enable spatially targeted reward delivery. **(B)** Overhead view of the gantry-mounted acrylic panel, which houses the reward dispensing assembly and prevents rats from climbing the gates. **(C)** Diagram of the reward dispensing mechanism, including a servo-actuated cannula, peristaltic pump, food reservoir, and tracking markers for real-time position alignment. Note that, before reward delivery, the right-angle elbow of the cannula is positioned over the center of a given chamber. In its default state, it is rotated out of reach of the rat through a slot in the acrylic panel. During reward delivery, the cannula tip is rotated downward to dispense reward directly onto the maze floor.

Reward is delivered via a stainless-steel cannula mounted to the gantry carriage (Fig. 6C). The cannula includes a 90° bend forming a pivoting arm actuated by a belt-driven micro-servo motor (SG90; TowerPro, Hong Kong, China), allowing it to rotate down to the maze floor during dispensing and retract back through a slot in the ceiling panel during transit. Liquid reward is delivered via a peristaltic pump (DFR0523; DFRobot, Shanghai, China), which is only activated when the cannula is lowered to prevent splashing or accidental dispensing on the animal. The mechanism is light and low-torque enough not to hurt the animal if the cannula lowers into it.

#### Motion Control

The gantry is controlled via GRBL firmware running on a dedicated Motion Control System (BlackBox X32, OpenBuilds, Zephyrhills, Florida, USA), which receives G-code jogging commands over a serial interface from a dedicated Arduino Mega 2560 (Fig. 2). The Arduino acts as an intermediary, receiving high-level homing, coordinate reset and movement ROS commands via EtherCAT and forwarding these to the GRBL controller.

#### ROS Integration

A dedicated ROS node governs gantry behavior and integrates real-time tracking data from both the rat and gantry from the tracking subsystem (Fig. 2). In continuous tracking mode, the gantry follows the rat using proportional control based on real-time rat and gantry pose data and halts when within 0.15 m of the target. Although the system supports this continuous animal tracking, in practice we opted for a chamber-level tracking policy. Under this configuration, the gantry remains stationary until the rat crosses into a new chamber, at which point it transitions to the corresponding chamber center. This approach was chosen to minimize behavioral disruption, as continuous gantry tracking was found to produce unwanted behavioral effects and occasional instabilities despite acceptable auditory noise levels.

The gantry control node also manages microcontroller and GRBL initialization, homing, mode transitions (e.g., idle vs tracking), reward triggering, cannula actuation, and pump activation. Reward delivery can be manually issued or initiated via a command that triggers a fixed actuation sequence: the cannula lowers, the pump activates for a user-defined time, and the cannula retracts. All low-level actuation is managed on the Arduino via two PWM channels (one for the servo, one for the pump). Gantry motion is temporarily disabled during reward delivery to ensure positional stability.

### Tracking Subsystem

#### Motion Capture Hardware

Rat position and orientation are tracked in 3D using a commercial motion capture system (OptiTrack, NaturalPoint Inc., Corvallis, OR, USA) comprising twelve PrimeX 13 infrared cameras (PX13; Fig. 7A) and two Prime Color video cameras (PCC0001; Fig. 7B). Reflective markers in a rigid configuration affixed to either a harness worn by the rat (Fig. 7C) or the recording headstage (Fig. 7D) allow for high-precision tracking at 240 Hz. The same system is used to track the gantry for real-time alignment of reward delivery (Fig. 6C and 7E).

**Figure 7.**
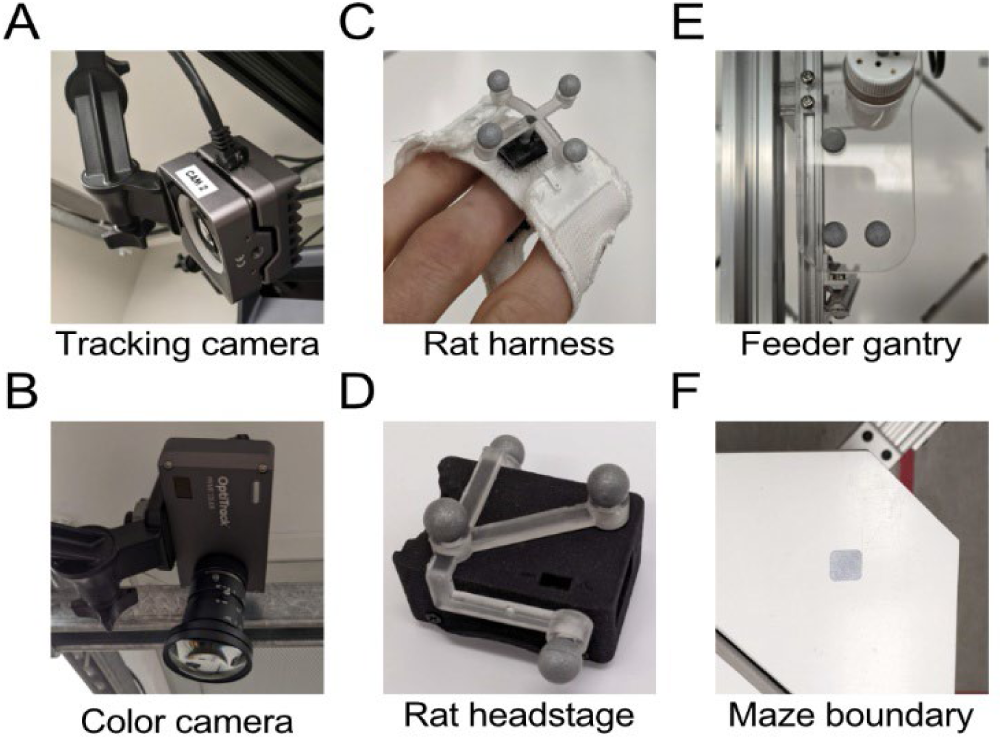
3D motion tracking components. **(A)** One of 12 IR cameras and **(B)** one of two color cameras used in the tracking system. Camera mounting positions are shown schematically in Figure 5A. Reflective markers are shown for tracking the rat via **(C)** a harness and **(D)** a recording headstage, as well as for tracking **(E)** the gantry and **(F)** the maze boundary. All elements are tracked in real-time at 240 Hz to enable closed-loop behavioral control.

Six reflective markers are permanently affixed along the maze perimeter and used to define the physical boundary and coordinate frame of the maze (Fig. 7F). Camera synchronization is managed via an OptiTrack eSync 2 synchronization module (OPTESY2), and data is streamed using the Motive software (version 3.3.2) in Tracker mode, which runs on the Windows computer.

#### ROS Integration

Raw tracking data for all elements (rat, gantry, and maze boundary) is streamed from Motive software to the Ubuntu ROS workspace via a custom ROS node that parses the data stream in OptiTrack NatNet protocol. A transformation between tracking world coordinate frame and the physical maze is computed from the known spatial configuration of three of the six reflective markers affixed along the maze perimeter, which define the maze-centered coordinate system. A maze transform node computes the transformations and continuously publishes poses for the rat’s harness / recording headstage and gantry within this reference frame. These poses are used throughout the ROS system to support closed-loop gate control, visual masking, gantry targeting, and other behavior-contingent processes. This setup enables low-latency, closed-loop behavioral control and targeted reward delivery.

### Graphical User Interface

A custom graphical user interface (GUI) was developed to enable interactive control, visualization, and configuration of the maze system during experimental sessions. The interface is implemented as a Qt-based plugin using PyQt5 and integrated into the ROS framework for live interaction with all subsystems.

#### System Configuration and Visualization

The main GUI window displays a real-time schematic of the maze, with each chamber and its surrounding gates rendered as clickable objects. Users can toggle the state of individual gates via mouse clicks. Gate configurations can be loaded from or saved to comma-separated value (CSV) files, and configurations can be browsed and previewed from the GUI. A color-coded status system provides diagnostic feedback for all gate modules, with gate state updates both driven by and published to ROS topics. Live rat position is rendered as an overlay on the schematic view.

System control buttons allow initialization, reinitialization, and shutdown sequences, including handshake verification with all connected Arduinos. User-editable system parameters include PWM duty cycle, movement timeouts, and maximum retries, each validated and transmitted at startup.

#### Stimulus and Reward Control

The GUI includes dedicated controls for gantry movement, rat tracking, cannula actuation, and pump operation. Reward delivery can be initiated manually or programmatically. Visual and auditory stimulus configurations can be sent to projectors using dedicated control buttons for gate wall panel and floor textures, defined in CSV files. Projector window alignment and full-screen toggling are also accessible via GUI buttons.

#### Experiment Session Control

A secondary GUI window provides session control and trial management. Users can load trial sequences from Excel files, preview or select individual trials, and step through the sequence manually or in order. The intention is that the Main GUI remains consistent across experiments whereas the experiment-specific GUI and controller can be customized by each experimenter. Trial definitions, including gate configurations and stimuli, are serialized and dispatched over ROS for real-time execution. Dedicated controls support session start, pause, and resume commands, as well as initiation of experiment-type-specific data recording using the rosbag format, including synchronized TTL pulse generation for external synchronization.

### Animals

All procedures were conducted in accordance with the Canadian Council on Animal Care guidelines and approved by the University of British Columbia Animal Care Committee. Two adult male Long-Evans rats were used in total.

Both rats were trained on the Contextual Conditional Discrimination task described below. Animals were housed on a 12-h light/dark cycle and tested during the light phase. All rats were food-restricted to approximately 80% of their free-feeding body weight and received liquid reward during testing.

For the electrophysiology experiments, one rat was implanted under isoflurane anesthesia with a 64-channel silicon probe (ASSY-236, Cambridge NeuroTech, Cambridge, UK) targeting dorsal CA1. The probe was secured with Metabond Quick (Parkell Inc., Edgewood, NY USA), dental cement and skull screws, and the animal was given 7 days to recover before recordings commenced. Neural signals were acquired with a SpikeGadgets HH128 Datalogger (SpikeGadgets, San Diego, CA, USA), sampled at 30 kHz to an onboard SD card. Eight non-functional channels were excluded, leaving 56 channels for analysis.

### Actuation Testing Protocols

The gate and gantry testing procedures described here were used for the gate timing, gantry movement, auditory noise, and electrophysiological noise tests. All tests were executed with custom Python scripts interfacing with the existing ROS nodes, which issued the movement commands and logged associated ROS topics.

#### Gate actuation protocol

The eight gates surrounding the center chamber were actuated in 20 cycles, each cycle consisting of one upward and one downward movement, yielding 40 movement events (Fig. 8A). Inter-movement intervals were fixed at 6 s. Gate movement commands and state messages (up/down) were logged to ROS bag files for analysis.

**Figure 8.**
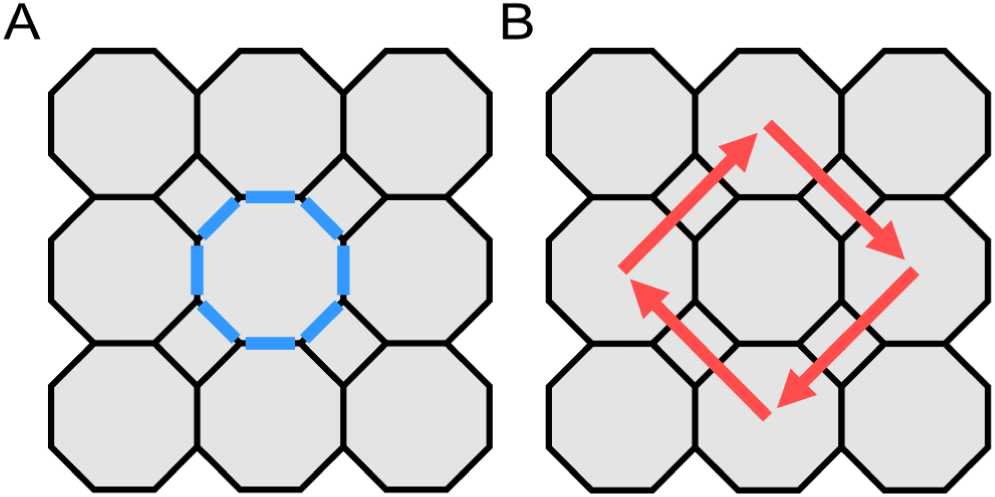
Gate and gantry actuation testing protocols. **(A)** The eight adjacent gates of the center chamber (blue) were repeatedly actuated in 20 full cycles (40 total up and down movements). **(B)** The gantry was moved between four outer chambers of the maze in a diamond pattern (red arrows), yielding a total of 20 movement events (5 per trajectory).

#### Gantry movement protocol

The gantry was moved sequentially between four outer chambers in a fixed diamond pattern, engaging both X and Y axes on each move (Fig. 8B). A total of 20 movement events were executed, evenly distributed across the four chamber-to-chamber transitions, with 5 s inter-movement intervals. Gantry movement commands and tracked pose data were recorded to ROS bag files for each transition.

### Gate and Stimulus Timing Test

Gate timing analysis was performed using the same procedure described above (Actuation Testing Protocols; Fig. 8). Gate latency was defined as the interval between the up or down move command message and the return message confirming actuation completion from the gate microcontroller.

For the stimulus timing, including image projection and audio playback from the four projectors, a second protocol was used in which projection and audio events alternated at fixed 5 s intervals. At the start of testing, the eight center chamber gates were raised. Two alternating sets of visual cues were projected onto the inner faces of the eight raised gates, yielding two images per projector and eight images in total. Alternating with this, a 1 s, 1 kHz auditory cue was played through all four projector-integrated speakers. A total of 40 events (20 projection, 20 audio) were presented per session. ROS messages from the command and feedback channels were logged for offline analysis. Projection latency was defined as the interval between the command message and the final render acknowledgment from all four projectors. Audio latency was defined as the interval between the command message and the onset of playback reported by the audio node.

### Gantry Movement Kinematic Test

We quantified gantry movement velocity, timing, and positional accuracy using the same test procedure as described above (Actuation Testing Protocols; Fig. 8). Motion capture data were aligned to commanded positions and, for each movement, arrival latency was defined as the time from command to the first frame in which the gantry position was within 2 cm of the target. Final accuracy was computed as the Euclidean distance between the target and the gantry’s position when it was nearest to the target. Speed profiles were obtained by windowing position traces for each move event, low-pass filtering (4th-order Butterworth, 6 Hz cutoff), and differentiating.

### Auditory Noise Test

To quantify the auditory levels generated by the Omniroute system’s mechanical components, sound recordings were collected during repeated movements of both the gate and gantry independently, as described above (Actuation Testing Protocols; Fig. 8). All recordings were taken from the center of the Omniroute maze using a high-fidelity ultrasound microphone (Model M500-384, Pettersson Elektronik AB, Uppsala, Sweden). Each event was preceded by a 1 kHz tone of 1 s duration, played through the projector-integrated speakers, which was used to align the audio recordings with the corresponding ROS event data during postprocessing. A portable 1 kHz sound-level calibrator (Model 8930, AZ Instrument Corp., Taichung, Taiwan) was used to record reference tones at 94, 104, and 114 dB SPL for offline calibration. Signals were sampled at 384 kHz and band-pass filtered from 200 Hz to 90 kHz.

Audio power was computed using a 20 ms moving RMS envelope applied to bandpass-filtered (200 Hz - 90 kHz), calibrated audio traces, then converted to decibels SPL using a single gain factor derived from reference tone recordings. For each event, a window was extracted beginning at movement onset. Window durations were 1.0 s for gate events and 1.5 s for gantry events. A corresponding baseline window of equal length was extracted 3.0 s (gate) or 2.5 s (gantry) after each movement event to avoid overlap with actuation noise.

For each window, mean SPL values were computed. Power spectral density (PSD) was estimated using Welch’s method (4096-sample Hann window, 50% overlap, 8192-point FFT). Signals were preprocessed identically to the SPL analysis and converted to Pascals prior to PSD computation. Condition-specific PSDs were averaged across events and smoothed using a 5-point moving mean for visualization, with results expressed in decibels relative to (20 µPa)²/Hz.

### Electrophysiological Noise Test

To assess potential electrical artifacts from gate and gantry movement, we recorded hippocampal neural signals during independent testing protocols for each subsystem described above (Actuation Testing Protocols; Fig. 8). For both tests, the animal was confined to a clear acrylic enclosure in the center chamber of the maze to restrict movement during recordings.

Continuous traces from 56 recorded channels were segmented into windows anchored directly to ROS movement onset times. The same windowing was used as the auditory noise test: 1.0 s for gate events and 1.5 s for gantry events, with matched baseline windows offset by 3.0 s (gate) or 2.5 s (gantry). For each window, signals were de-meaned per channel and RMS amplitude was computed per channel, then averaged across channels. PSD was estimated using Welch’s method (4096-sample Hann window, 50% overlap, 8192-point FFT), averaged across channels and events, and expressed in decibels relative to 1 µV²/Hz.

### Contextual Conditional Discrimination Task

Animals were trained to perform a novel Contextual Conditional Discrimination task using the Omniroute system. The maze was configured as a T-maze with one start chamber, a choice chamber, and two possible goal chambers (Fig. 9). The task relied on two complementary types of visual cues. A floor-projected “start-cue” (green light covering the maze floor) signaled whether the rat should approach the visually marked or the unmarked chamber, which was designated by the “goal-cue” (green triangles projected onto the interior faces of the gate wall panels for one goal chamber). Goal-cue location was randomized across trials, such that the rat had to interpret the start-cue to determine whether to approach the triangle-marked chamber (goal-cue) or the unmarked (dark) chamber. The two “cue-rule” contingencies were: green floor → go to the green-triangle chamber; no floor projection → go to the dark chamber.

**Figure 9.**
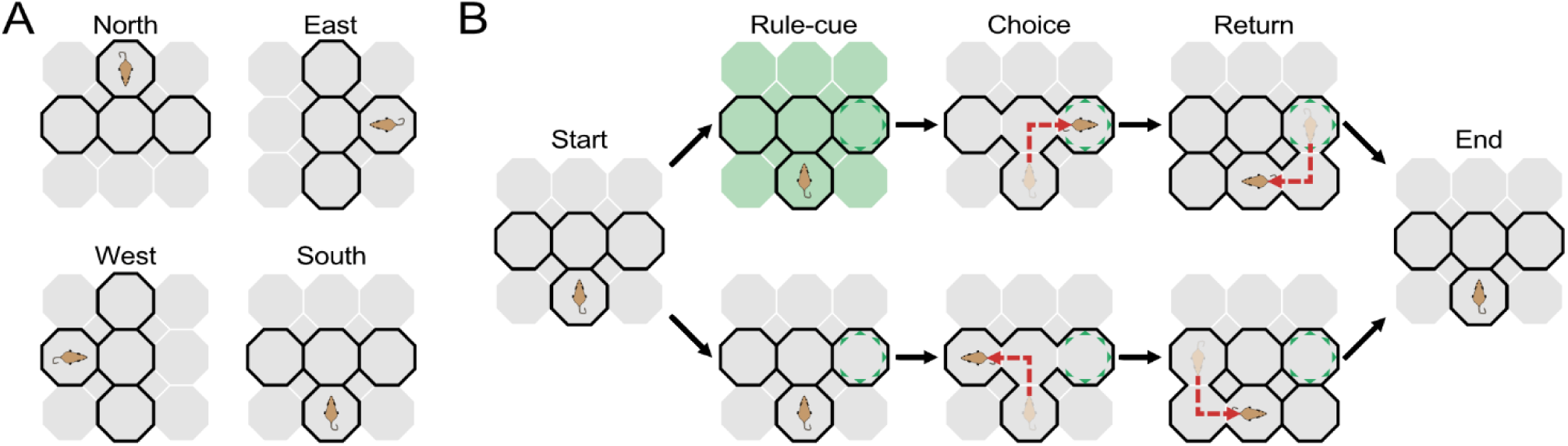
Task overview for the Contextual Conditional Discrimination task. **(A)** For each trial, the maze was configured as a T-maze in one of four possible orientations (North, East, West, South), as defined by the location of the start chamber. This design preserved the relative T-maze geometry while varying the spatial positions of the start, choice, and goal chambers. **(B)** Example trial sequences with the animal beginning in the south start chamber and the goal-cue located in the right goal chamber. In practice, the goal-cue location varied across trials between the left and right goal chambers. Red dashed paths show the rat’s trajectory from the start chamber through the choice chamber to the correct goal chamber. Green triangles indicate the goal-cue projected onto the interior gate wall panel faces of the assigned goal chamber. Top row: with the start-cue present (green floor projection), the correct choice is the triangle-marked (goal-cue denoted) goal chamber. Bottom row: with the start-cue absent (no floor projection), the correct choice is the unmarked (non-goal-cue denoted) goal chamber. Trial sequence: rats began confined in the start chamber, where the start-cue was either presented or absent. After a fixed delay, gates to the choice and goal chambers opened, allowing the rat to navigate to its selected goal. Upon entry, the goal chamber gate closed and reward was delivered if the choice matched the current cue-rule contingency. After a delay, gates opened to allow the rat to return to the same start chamber for the next trial or to route it to a new start chamber (not shown).

Each trial began with the rat confined in the start chamber. The start-cue was presented on the floor for a brief period, followed by a short delay. The gates to the choice and goal chambers were then opened, allowing the rat to select a goal. Upon entry into a goal chamber, the gate was closed and reward was delivered if the choice matched the current cue-rule contingency. After a delay, alternate gates were opened to allow the rat to return to the start chamber via side chambers. All gate movements, cue presentations, and rat position data were recorded automatically for offline analysis.

Training proceeded in three phases. In Phase 1, rats were trained on a single cue-rule contingency until achieving n consecutive correct trials, with n reduced across sessions (10, 8, 6, 4, 2). Once criterion was met, the active cue-rule contingency switched (e.g., from “green floor → triangle chamber” to “dark floor → dark chamber”). This block-based shaping established the cue-choice association. In Phase 2, the active cue-rule contingency was randomized on each trial, requiring flexible use of the start-cue rather than relying on block structure. In Phase 3, the entire T-maze layout was rotated counter-clockwise among four orientations after blocks of 10-20 trials, altering the spatial locations of the start and goal chambers while leaving the cue-rule contingencies unchanged. This manipulation dissociated cue-based decision-making from fixed spatial trajectories.

It is important to note that the gantry system was not utilized for any of the behavioral data presented. Instead, liquid food rewards consisting of equal parts water and chocolate Ensure meal replacement drink (Abbott Laboratories, Chicago, IL, USA) were manually delivered via pipette within the respective goal chamber.

### Statistical Analysis and Data Processing

Data preprocessing was performed in Python 3 to extract and align ROS and tracking data from raw recordings. All subsequent analyses and plotting were conducted in MATLAB 2022 (MathWorks, Natick, MA). Unless otherwise noted, statistical tests were two-tailed with α = 0.05. Results are reported as mean ± SD with n indicated, and exact p-values are given to three significant figures when p ≥ 0.001 or as p < 0.001 when smaller. Error bars represent SD unless otherwise noted.

### Design and Code Repository Access

CAD models and laser- and waterjet-cutting templates for the projector mounts, gantry, and maze platform and frame of the Omniroute system are publicly available through the Open Science Framework (OSF) page (https://osf.io/xfpsv/). For the gate modules and associated custom motor driver and Cypress I/O expander PCBs, more detailed design files, documentation, and CAD models are available through a sperate OSF repository (https://osf.io/uy7ez).

The control codebase is divided across two GitHub repositories, corresponding to the two ROS workspaces used in the system architecture (Fig. 2):

- Ubuntu: https://github.com/NC4Lab/omniroute_ubuntu_ws/tree/lester2025omniroute
- Windows: https://github.com/NC4Lab/omniroute_windows_ws/tree/lester2025omniroute

All resources are released under the CERN Open Hardware License – Weakly Reciprocal (CERN-OHL-W) and are intended to facilitate reuse, modification, and extension by the research community.

## Results

### Gate and Stimulus Timing

To assess command latencies within the system, we measured the round-trip time between command messages and subsystem feedback acknowledgments for gate actuation, audio playback, and projected image rendering. For the gate tests, we used the protocol described above (Actuation Testing Protocols; Fig. 8). Across all events, mean gate actuation latency was 915.5 ± 47.9 ms (n = 40) (Fig. 10A). Upward movements were consistently slower than downward movements, averaging 959.7 ± 23.2 ms and 871.3 ± 7.5 ms, respectively (paired two-tailed t test: t(19) = 14.96, p = 5.787e-12, Cohen’s d = 3.34, 95% CI [76.05, 100.79]; n = 20 pairs). Projection render latencies were substantially lower in magnitude and variability (Fig. 10B). The average latency was 358.7 ± 12.8 ms across 20 events, with individual latencies tightly clustered around the mean. Audio playback latencies were intermediate between projection and gate actuation (Fig. 10C), with playback beginning after an average delay of 724.6 ± 7.2 ms (n = 20) and exhibited minimal variability across events.

**Figure 10.**
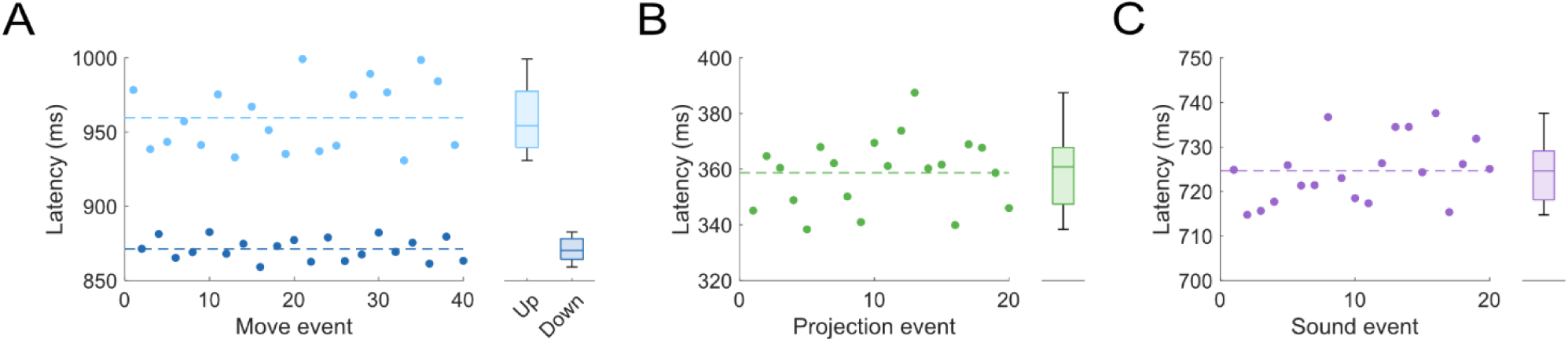
Gate actuation and stimulus presentation timing. Latencies were measured between ROS command messages and feedback acknowledgements for gate events as well as projection and audio stimulus delivery. **(A)** (left) Latency for gate up (light blue) and down (dark blue) movement events and mean latency of each movement direction (dashed lines). (right) latency distribution across all events with the median and interquartile range indicated (n = 40). The mean latency was 959.7 ± 23.2 ms for up and 871.3 ± 7.5 ms for down movements. **(B)** Projection render latencies shown using the same format as A for each projection event (n = 20), with a mean latency of 358.7 ± 12.8 ms. **(C)** Audio playback latencies shown using the same format as A for each audio event (n = 20), with mean latency of 724.6 ± 7.2 ms.

Together, these results show that stimulus presentation events (projection and audio) and gate actuation occur with reasonably consistent sub-second timing, although gate actuation introduces longer delays that differ systematically by movement direction.

### Gantry Movement

To evaluate the performance of the reward delivery subsystem, we characterized the movement and positioning accuracy of the gantry using the protocols described above (Actuation Testing Protocols; Fig. 8), in which the gantry was jogged approximately 0.43 m per trajectory in a diamond pattern between four outer chambers of the maze, engaging both X and Y drive axes for each translation.

The measured gantry position closely tracked commanded positions along both axes, with minimal observable overshooting or oscillation at the end of each move (Fig. 11A). Speed profiles showed smooth, symmetric acceleration and deceleration patterns and minimal variability across the four movement trajectories (Fig. 11B), with peak speed 0.474 ± 0.001 m/s and mean speed 0.210 ± 0.001 m/s.

**Figure 11.**
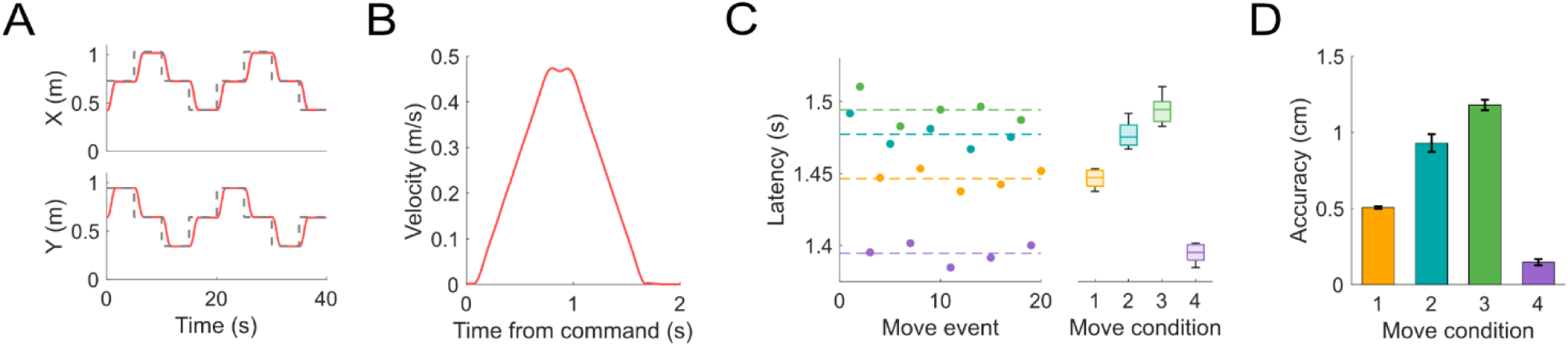
Gantry movement kinematics. The gantry was sequentially translated between four outer chambers in a fixed diamond pattern, same trajectories as in Figure 8B, covering approximately 0.43 m for each traversal. **(A)** Example time-series showing commanded positions (dashed) and measured gantry pose (solid) along the X and Y axes, for the first two full cycles of movement. **(B)** Mean ± SD velocity profile across all movements, aligned to command onset, derived from low-pass-filtered position traces. Velocity profiles showed minimal variability between or within move conditions. **(C)** Arrival latencies are shown as individual points color-coded by the four movement trajectory conditions (orange, teal, green, purple) with dashed lines indicating the per-condition mean (left panel), and as distributions using a box-and-whisker style chart with the median and interquartile range indicated (right panel). **(D)** Final position accuracy by move condition. Across movements, mean arrival latency was 1.45 ± 0.04 s and final accuracy was 0.69 ± 0.41 cm.

Across all moves, the latency between when move commands were sent and when the gantry reached its target was 1.45 ± 0.04 s. Within-condition variability in arrival latency was even lower, with standard deviations ranging between ∼30-50 ms. Between movement conditions, the mean arrival latency differed by ∼100 ms, reflecting modest trajectory-specific effects (Fig. 11C).

The average positional accuracy was 0.69 ± 0.41 cm across all moves. Within-condition variability was minimal, with standard deviations of ∼0.1-0.6 cm depending on condition. Between conditions, mean accuracy differed by ∼1.1 cm, again suggesting trajectory-specific effects. Accuracy remained high across all conditions, with all targets reached well within the 2 cm tolerance (Fig. 11D).

### Gate and Gantry Auditory Noise

We assessed the acoustic profile of gate and gantry movements under typical operating conditions using high-resolution acoustic recordings taken from the center of the Omniroute maze during the testing procedure described (Actuation Testing Protocols; Fig. 8). Sound pressure levels and spectral content were then evaluated and compared between active and baseline periods.

Gate actuation produced transient increases in broadband sound pressure levels relative to baseline (Fig. 12A). Across 20 cycles of upward and downward gate movement, mean SPL rose from a baseline of 76.19 ± 1.41 dB SPL to 102.83 ± 1.16 dB SPL during movement windows (Fig. 12B). After baseline correction, the average increase in mean ΔSPL was +26.64 ± 1.16 dB. Power spectral density analysis revealed a broadband elevation during movement periods without the emergence of narrowband peaks or frequency-specific artifacts (Fig. 12C).

**Figure 12.**
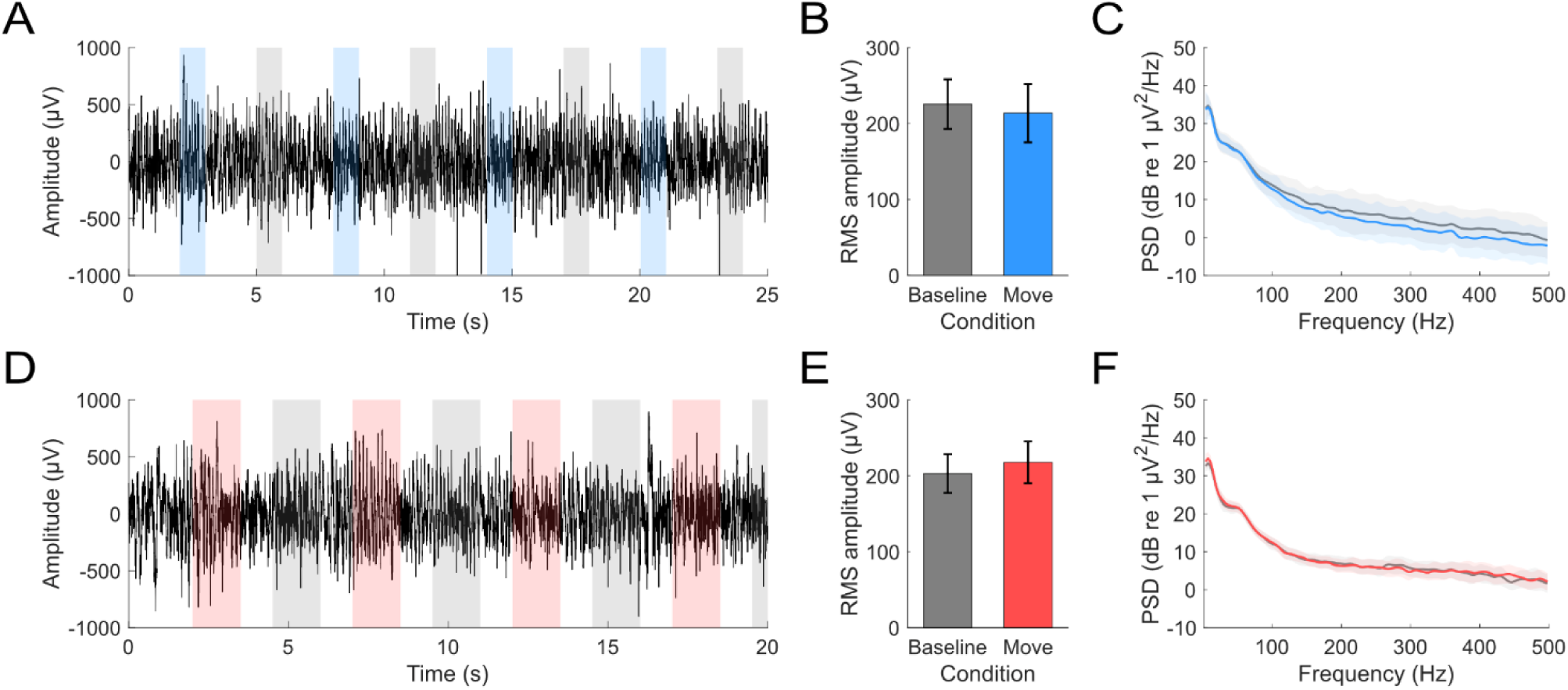
Gate and gantry auditory noise characteristics. Recorded sound pressure levels from the center of the maze were compared between actuation and matched baseline periods without actuation. **(A-C)** Gate testing: **(A)** Baseline-corrected ΔSPL waveform across the first 4 of 40 movement events, with movement windows in blue and baseline windows in gray. The average baseline-corrected SPL during gate movement was 26.64 ± 1.16 dB. **(B)** Mean sound pressure level (dB SPL) during baseline and gate movement periods (mean ± SD). **(C)** PSD of the calibrated waveform across conditions (dB re (20 µPa)²/Hz), showing a broadband elevation during movement without narrowband features. **(D-F)** Gantry testing: **(D)** ΔSPL waveform with movement windows in red and baseline windows in gray, shown for the first 4 of 20 events. The average baseline-corrected SPL during gantry movement was 19.56 ± 0.89 dB. **(E)** Mean SPL by condition (mean ± SD). **(F)** PSD comparison shows a similar broadband elevation during gantry motion, without dominant peaks, though minor frequency-specific elevations are present.

Movement of the gantry also resulted in transient increases in sound pressure level, though to a lesser extent than gate actuation (Fig. 12D). During 20 gantry translation events, mean SPL increased from a baseline of 76.72 ± 0.38 dB SPL to 96.28 ± 0.89 dB SPL during movement windows (Fig. 12E), resulting in a baseline-corrected average increase of +19.56 ± 0.89 dB. Spectral analysis showed a broadband elevation across the full frequency range during gantry motion, with two notable discrete spectral peaks observed at ∼2 kHz and ∼53 kHz (Fig. 12F).

Overall, these findings indicate that auditory noise introduced by gantry movement was lower in amplitude than that produced by gate actuation. Both systems generated broadband increases in sound pressure level, but mean baseline-corrected SPL elevations remained below ∼20 dB for gantry movements and below ∼30 dB for gate movements. Gate movement did not introduce dominant tonal artifacts, though the PSD from gantry movements did suggest possible frequency-specific artifacts.

### Gate and Gantry Electrophysiological Noise

We evaluated whether gate and gantry actuation introduced electrical artifacts into neural recordings from a chronically implanted rat confined to the center chamber of the Omniroute maze. Analyses were performed on 56 of 64 recorded channels placed in dorsal CA1 after exclusion of non-functional channels. Both the gate and gantry tests followed the same protocol described above (Actuation Testing Protocols; Fig. 8), and were conducted with the rat confined to the center chamber of the maze in a clear acrylic enclosure.

For the gate test, the average RMS amplitude computed across channels showed no significant difference between movement and baseline periods (baseline: 225.75 ± 32.61 µV; move: 213.82 ± 38.46 µV; paired two-tailed t test: t(39) = 1.93, p = 0.061, Cohen’s d = −0.30, 95% CI [−0.60, 24.47]; Fig. 13A and B). Spectral analysis similarly revealed overlapping power spectra across movement and baseline conditions, with no frequency-specific deviations observed across the 1-500 Hz range (Fig. 13C).

**Figure 13.**
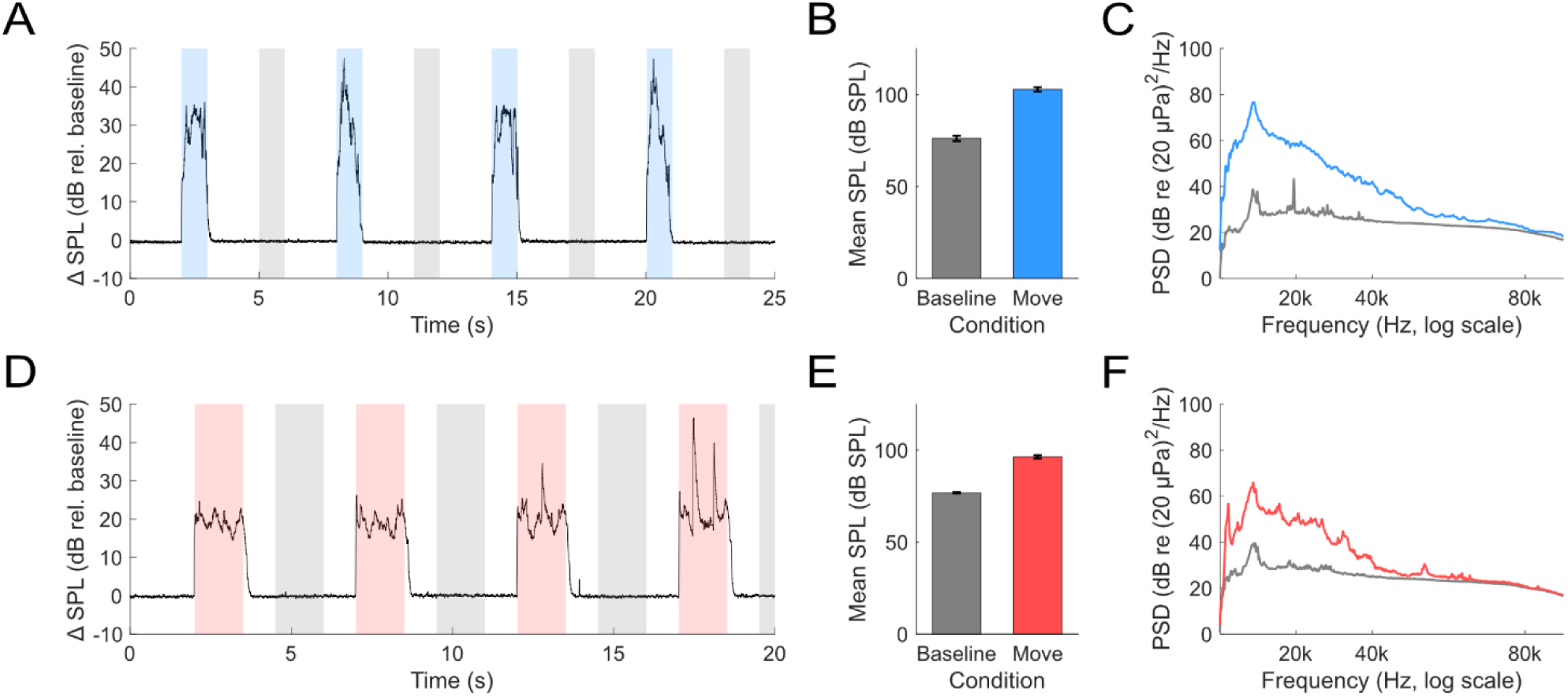
Gate and gantry electrophysiological noise characteristics. Continuous neural signals from 56 of 64 recorded channels in the hippocampal CA1 region in a chronically implanted rat during independent movement of each of the gate and gantry subsystems. **(A-C)** Gate testing: The eight gates surrounding the center chamber were actuated in 20 full cycles (40 total movement events; Fig. 8A). **(A)** Average voltage trace across 56 channels during the first 4 of 40 movement events. Shaded windows indicate movement periods (blue) and matched baseline periods (gray). **(B)** RMS amplitude computed per channel after de-meaning and averaged across channels during movement and baseline windows (mean ± SD). **(C)** Mean PSD during gate movement and baseline windows (mean ± SD), expressed in dB re 1 µV²/Hz. Spectral profiles overlapped across conditions, with no frequency-specific deviations observed. **(D-F)** Gantry testing: Recordings were obtained during 20 movement events as the gantry traversed a diamond path across the outer chambers (Fig. 8B). **(D)** Average voltage trace across the first 4 of 20 events, with movement (red) and baseline (gray) windows indicated. **(E)** RMS amplitude during gantry movement and baseline windows (mean ± SD). **(F)** Mean PSD during gantry movement and baseline periods (mean ± SD). As in the gate condition, no frequency-specific changes were observed during gantry motion.

For the gantry test, no significant change in mean RMS amplitude was observed between movement and baseline windows (baseline: 202.97 ± 25.51 µV; move: 217.82 ± 27.52 µV; paired two-tailed t test: t(19) = −1.74, p = 0.098, Cohen’s d = 0.39, 95% CI [−32.69, 3.00]; Fig. 13D and E). PSD profiles during gantry movement closely matched those observed during baseline, with no emergence of frequency-specific artifacts (Fig. 13F).

Together, these results indicate that neither gate nor gantry movement introduces detectable electrical artifacts into neural recordings under typical operating conditions.

### Task Performance

To assess how the Omniroute performs in real-world behavioral experiments, two rats were trained on our Contextual Conditional Discrimination task. The task required animals to flexibly combine two sources of visual information to guide goal selection. Success depended on interpreting the start-cue (floor-projected green light) at the start of the trial to determine whether to select the goal-cue chamber (the chamber with green triangles projected on internal gate wall panel faces) or the unmarked chamber, and applying this rule consistently across changing spatial orientations of the maze (Fig. 14).

**Figure 14.**
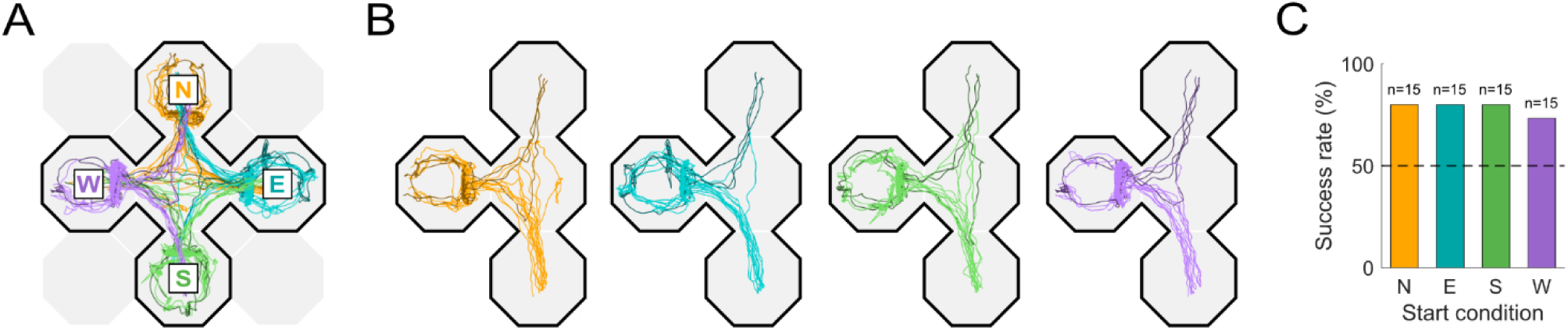
Example Phase 3 trajectories and performance on the Contextual Conditional Discrimination task. **(A)** Raw trajectories from a single Phase 3 session (Rat 1). Each line shows the rat’s path from one of four start chambers (N, E, S, W), color-coded by start location (orange, teal, green, purple). Correct trials are plotted with brighter lines and errors with darker lines. **(B)** Normalized trajectories from the same session, plotted separately by start location. For each panel, trajectories were rotated to a common orientation and mirrored into a shared reference frame such that the correct goal chamber appears on the right for all trials. **(C)** Success rate for this session by start location (n = 15 trials per start), computed as the percentage of correct trials out of total trials. Overall performance was 47 / 60 correct trials (78%), with success rates ranging from 73-80% across the four start conditions.

We first analyzed trajectories from a representative Phase 3 session. In this phase, the cue-rule contingency (presence or absence of the start-cue) and the location of the goal-cue varied across trials, while the maze orientation (the entire T-maze rotated among four start positions: North, East, South, and West) varied across blocks of trials. The rat exhibited reliable navigation performance across all start conditions (Fig. 14). Raw trajectories overlaid on maze coordinates showed consistent convergence to the correct goal chambers from each of the four start positions, with errors diverging toward the opposite chamber (Fig. 14A). When normalized to a canonical reference frame, trajectories from each start condition overlapped closely, with most trial trajectories targeted to the correct goal chamber, highlighting that the rat applied the cue-based rule consistently rather than relying on fixed spatial paths (Fig. 14B). Success rate was computed for each start orientation block as the percentage of correct goal choices. Success rates were uniformly high across start positions, ranging from 73-80% correct (overall 47/60 trials, 78%; Fig. 14C). These results demonstrate that the rat generalized the cue-based decision rule across different maze orientations, flexibly integrating the rule- and goal-cues to guide navigation.

Behavioral performance was assessed for both rats across sessions and training phases based on their success rates when performing the task (Fig. 15). Across sessions, both rats showed a clear acquisition of the task in Phase 1 and maintained above-chance accuracy thereafter (Fig. 15A and B).

**Figure 15.**
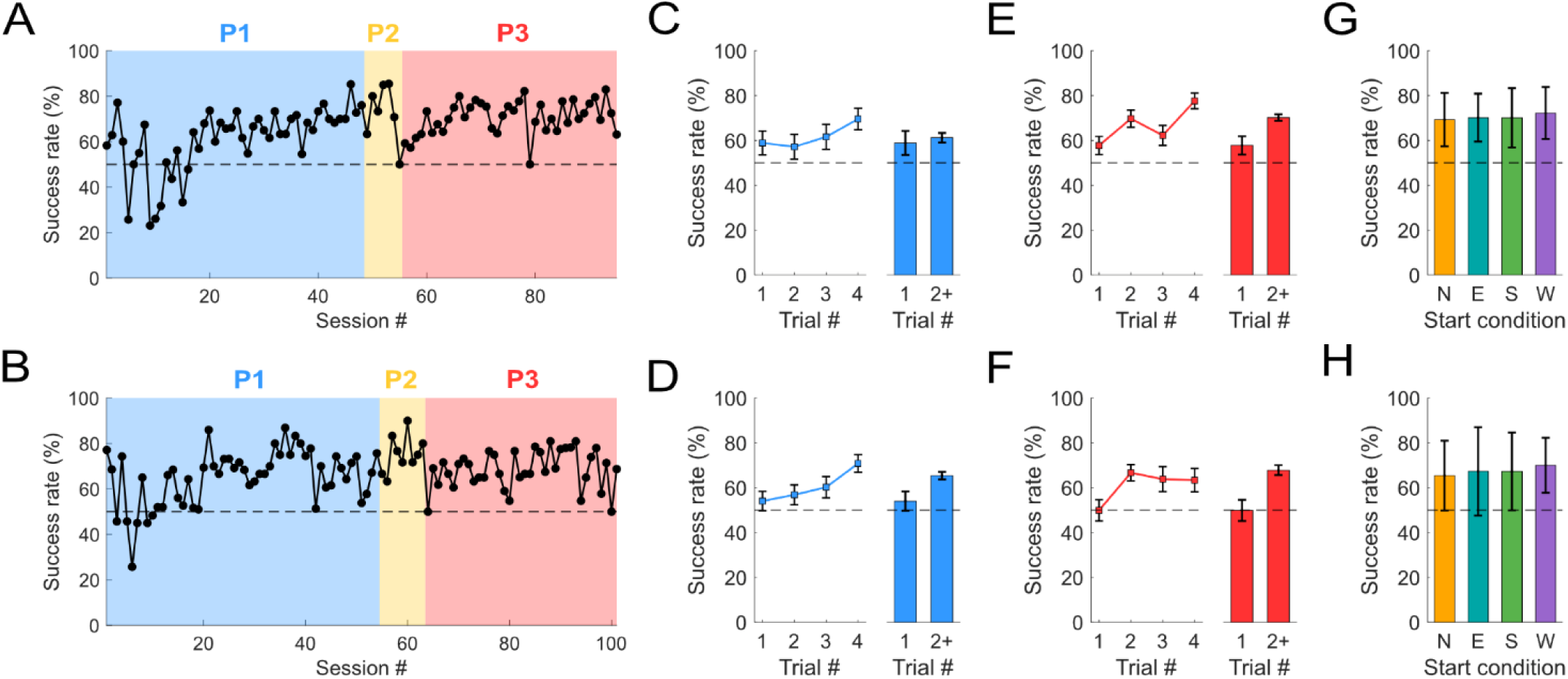
Aggregate performance across training phases on the Contextual Conditional Discrimination task. **(A-B)** Session-level success rates, computed as the percentage of correct trials out of total trials, for Rat 1 **(A)** and Rat 2 **(B)**. Colored background patches indicate training Phases 1-3 (blue, yellow, red). Both rats exhibited reliable acquisition in Phase 1 and maintained above-chance performance across subsequent phases. **(C-D)** Trial-level performance in Phase 1, aligned to the first trial after the cue-rule contingency switched (e.g., green floor projection to no floor projection or vice versa). Line plots show mean ± SEM success rates across the first four trials following each switch, averaged across sessions (left). Bar plots compare performance on the first trial versus the mean of all subsequent trials (2+; right) for Rat 1 **(C)** and Rat 2 **(D)**. Both rats tended to perform worse on the first trial after a rule change, with improved accuracy on subsequent trials, though this difference was only significant for Rat 2. **(E-F)** Trial-level performance in Phase 3, aligned to the first trial after the maze orientation was rotated. Plots are organized as in **(C-D)**. Both rats performed significantly worse on the first trial of each new orientation compared to later trials. **(G-H)** Success rates in Phase 3 broken down by start location (N, E, S, W) for Rat 1 **(G)** and Rat 2 **(H)** color-coded by start location (orange, teal, green, purple). Bars show mean success rates across sessions for each start orientation (mean ± SD across sessions), with colors corresponding to the four start positions (orange, teal, green, purple). Both rats maintained consistently high success across all start orientations.

To quantify initial acquisition, success rates were compared between the first and last half of Phase 1 training sessions. Rat 1 showed a significant increase between the halves (first half: 53.8% ± 13.6; last half: 68.5% ± 9.2; independent two-sample t test: t(31) = −4.20, p < 0.001, Cohen’s d = 1.21, 95% CI [−21.85, −7.56]), as did Rat 2 (first half: 60.7% ± 11.8; last half: 69.9% ± 10.4; independent two-sample t test: t(44) = −2.96, p = 0.005, d = 0.81, 95% CI [−15.34, −2.92]). These findings indicate that both rats acquired the task during Phase 1.

Session-level performance was then compared to chance (50%) using one-sample t tests. Both rats performed significantly above chance in Phase 1 (Rat 1: 61.2% ± 14.1, t(47) = 5.49, p < 0.001, d = 0.79, 95% CI [57.09, 65.29]; Rat 2: 65.1% ± 12.1, t(53) = 9.14, p < 0.001, d = 1.24, 95% CI [61.78, 68.40]), Phase 2 (Rat 1: 72.6% ± 12.7, t(6) = 4.69, p = 0.003, d = 1.77, 95% CI [60.78, 84.35]; Rat 2: 75.4% ± 8.3, t(8) = 9.19, p < 0.001, d = 3.06, 95% CI [69.01, 81.74]), and Phase 3 (Rat 1: 70.7% ± 7.3, t(39) = 18.04, p < 0.001, d = 2.85, 95% CI [68.39, 73.04]; Rat 2: 68.5% ± 8.3, t(37) = 13.74, p < 0.001, d = 2.23, 95% CI [65.76, 71.22]). These results confirm that both rats acquired the cue-based decision rule during Phase 1 and sustained reliable above-chance performance in subsequent phases.

To determine how rats adjusted to contextual cue changes during Phase 1, when the start-cue was systematically varied in blocks of 10-20 trials (e.g., green floor projection to no floor projection), we analyzed success rates on the first trial following each cue switch relative to the average across later trials (Fig. 15C and D). Rat 1 showed no significant difference (first trial: 61.3% ± 15.7; later trials: 63.6% ± 14.4; paired two-tailed t test: t(47) = −0.40, p = 0.689, d = 0.06, 95% CI [−14.10, 9.39]), whereas Rat 2 showed significantly lower performance on the first trial compared to subsequent trials (first trial: 59.4% ± 13.2; later trials: 70.8% ± 12.1; paired two-tailed t test: t(53) = - 2.67, p = 0.010, d = 0.36, 95% CI [−19.89, −2.81]). These results suggest that while both animals required some acclimation to start-cue changes, only Rat 2 exhibited a consistent decrement on the first trial of a new block.

Similarly, we assessed the effect of maze orientation changes in Phase 3 by aligning performance to the first trial after each rotation of the T-maze (Fig. 15E and F). Both rats showed significantly lower success rates on the first trial following a rotation compared to the average across later trials (Rat 1: first trial: 58.2% ± 10.1; later trials: 70.7% ± 7.4; paired two-tailed t test: t(39) = −2.91, p = 0.006, d = 0.46, 95% CI [−21.10, −3.81]; Rat 2: first trial: 50.6% ± 11.4; later trials: 68.5% ± 8.4; paired two-tailed t test: t(37) = −3.94, p < 0.001, d = 0.64, 95% CI [−27.12, −8.70]), indicating a brief orientation-specific adjustment period before consistent application of the cue-guided rule. We also compared success rates across the four start orientations in Phase 3 by averaging performance across sessions for both rats (Fig. 15G and H). Both rats performed similarly from each start location, and no consistent differences in performance were observed between orientations, again suggesting that application of the cue-guided rule generalized across all start conditions.

## Discussion

The Omniroute system supports real-time behavioral adaptation and precise temporal coordination between stimulus events, physical environment geometry, and reward delivery. All subsystems are designed to operate programmatically and in closed-loop with the animal’s behavior. The system architecture and constituent hardware elements are implemented using affordable, open-source components, except for the proprietary tracking system. The full codebase and design files are available through our OSF page and associated GitHub repositories to support replication and adaptation. The gate modules, custom motor driver boards, and Cypress I/O expander interface have been released previously through a separate OSF repository and are described in detail in prior work (Lester et al., 2024). System validation confirmed that execution timing across subsystems was stable and within a sub-second range, with reproducible round-trip latencies for gate, projection, and audio events. The gantry achieved accurate XY positioning with centimeter-level precision. Actuation of both gates and gantry elevated sound pressure levels in a broadband fashion but with minimal narrowband spectral features, and neither produced measurable artifacts in local field potential recordings. Finally, behavioral testing demonstrated that rats could acquire and perform a contextual conditional discrimination task within the system, confirming its utility for flexible, cue-guided navigation experiments utilizing dynamically configurable routes.

The gate subsystem behaved predictably and at time scales sufficient to reliably control animal access and progression through the maze. In the integrated system, command-to-completion latencies were consistently sub-second (915.5 ± 47.9 ms), though upward movement (∼960 ms) was systematically slower than downward (∼870 ms). These latencies reflect the full control path from ROS command to completion acknowledgment, which is what should be considered for coordinating closed-loop scheduling. As we reported previously, the actual mechanical travel time for our gate modules was 578 ± 17 ms for upward movements and 531 ± 13 ms for downward movements when testing one of five modules at a time on a stand-alone system (Lester et al., 2024). Thus, faster instantiation can be realized if movement is scheduled directly from the microcontroller to account for the round-trip command times. Even these latencies, however, are likely not responsive enough to redirect a fast-approaching rat, so experiments must be structured such that gates are positioned sufficiently ahead of an animal’s estimated arrival.

System-timing tests for visual and auditory stimulus presentation also showed command-to-completion latencies in the sub-second range, with projected images rendering in ∼359 ± 13 ms with minimal variability and audio playback beginning within ∼725 ± 7 ms with equally low variability. While these latencies are acceptable for most trial-level and some closed-loop paradigms, including our own, they could be improved with straightforward modifications to the codebase, such as pre-queuing stimuli based on known render and playback latencies or maintaining persistent output streams for audio playback rather than reinitializing devices on each use. For experiments requiring tighter precision for image display, further gains are possible through OpenGL optimization (e.g., pre-loading images into GPU memory, minimizing texture sizes and shifting homography calculations to shaders). In the current implementation, however, both subsystems operate within timescales that are sufficient for trial-based cue delivery and many applications requiring behavior-contingent stimulus delivery.

Tests of gantry movement confirmed predictable chamber-to-chamber translations with smooth velocity profiles and high precision targeting. For these tests, the gantry traversed a fixed diamond pattern between outer chambers, covering approximately 0.43 m per move. Variability in travel velocity (mean: 0.210 ± 0.001 m/s; peak: 0.474 ± 0.001 m/s) and total travel time (1.45 ± 0.04 s) was low across trajectories, and final position error remained sub-centimeter (0.69 ± 0.41 cm). These properties are more than sufficient for spatially discrete, pre-specified reward delivery, such as the chamber-centered policy used here, although trajectory-specific biases in accuracy likely reflect minor pose-alignment errors in the tracking-to-maze transform.

While the system supports continuous real-time animal tracking, in practice, we chose to use a chamber-to-chamber policy, as small corrective motions during continuous tracking proved aversive for some animals. This approach still allows the gantry to act as a barrier to climbing and to provide spatially and temporally precise reward delivery. If required, these limitations could be mitigated using predictive tracking methods, such as Kalman filtering or model predictive control, to smooth trajectories and anticipate rapid changes in position. It is also worth noting that the use of a commercial 3D tracking system like the one we use is not required, even for real-time animal tracking. In practice, any solution that supports stable transformation of 2D position to real-world coordinates at an adequate frame-rate and low latency is sufficient for the required closed-loop control of our system (Mathis et al., 2018; Pereira et al., 2022).

Acoustic and electrical noise tests confirmed that both gate and gantry actuation produce short-lived, broadband acoustic transients without introducing detectable electrical artifacts into hippocampal recordings. For acoustic measures, broadband sound pressure levels increased by roughly 25-30 dB above baseline for gate movements and ∼20 dB for gantry translations. These brief, broadband changes appear comparable to the transient sound increases that would commonly be encountered in standard housing environments. Gate actuation did not introduce any evident narrowband tonal peaks, though discrete spectral peaks were observed at ∼2 kHz and ∼53 kHz during feeder gantry traversals. Interference with frequency-specific stimuli should still be negligible if cue delivery is restricted to non-motion periods or confined to frequencies outside this range. Electrophysiological recordings taken from a chronically implanted animal likewise showed minimal evidence of movement-locked artifacts for either subsystem across the 1-500 Hz range, confirming that both mechanical subsystems can be used concurrently with neural data collection without introducing measurable electrical contamination.

To demonstrate the Omniroute system’s utility under the kinds of applications that highlight its strengths, we tested freely moving rats on our Contextual Conditional Discrimination task. This task required animals to integrate two temporally segregated visual cues to guide decision-making within a T-maze configuration of the Omniroute maze. During this task, both the goal location and the orientation of the maze were varied across trials, resulting in a paradigm that required flexible decision-making rather than reliance on fixed spatial geometries or relationships.

Both animals acquired the task and performed significantly above chance across all three phases of training. Most of the learning occurred during Phase 1 of training. Example session data showed stereotyped running trajectories that largely converged on the correct goal from all four start positions, and aggregate analysis confirmed comparable performance across start conditions for both rats. Trial-aligned analyses showed small first-trial costs after task changes: accuracy dipped significantly on the first trial after rule switches for one rat compared to later trials, and both rats showed a significant first-trial decrement after maze rotation. Taken together, these findings show that rats could acquire and flexibly apply the rule rather than rely on a static route, even when the maze orientation was varied.

Two qualifications are important to note for this component of the study. First, the behavioral dataset is limited to two animals, so estimates of effect size and generality are provisional. Second, rewards in these experiments were delivered manually; the gantry was validated kinematically but not used during the behavioral testing reported here. While these animals were being trained, the gantry system was not fully operational. The Omniroute system can, nonetheless, support automated reward delivery and closed-loop policies, and ongoing work is underway that incorporates the feeder gantry into similar tasks. Together, these results demonstrate that the dynamically configurable stimuli and route functionality of Omniroute can be used to test the flexibility of cue-guided navigation across changing spatial contexts in freely behaving rats, while highlighting straightforward next steps for scaling and full system integration.

To our knowledge, no single existing platform simultaneously combines continuous 2D locomotion with real-time control over routes, multimodal cues, and reward delivery, as has been validated with our Omniroute system. Many of the existing automated platforms are proprietary with limited versatility, such as the Multi-Maze system (Multi-Maze; Ugo Basile Srl, Italy) or the Automated Maze Suite (MazeSuite; ConductScience, USA). Even open, reconfigurable maze designs are typically restricted in the types of configurations supported and lack the design flexibility to support rapid and programmatically reconfigurable spatial contingencies under closed-loop control (Holleman et al., 2019; Sawatani et al., 2022; Jankowski et al., 2023; Porter et al., 2025). One notable exception is the “honeycomb maze” (Wood et al., 2018), which, while highly reconfigurable, uses a series of elevated platforms that prevent continuous running and may induce anxiety in rodents.

Head-fixed VR systems now allow for precise control over environment geometry, sensory inputs, and food-based reinforcement beyond what is achievable with real-world navigation paradigms. These benefits, however, come at the cost of natural motor engagement. Head-fixed rodents navigating VR environments exhibit altered network dynamics, including reduced theta modulation, disrupted entorhinal grid coding, and degraded hippocampal spatial tuning relative to free behavior (Chen et al., 2013; Ravassard et al., 2013; Aghajan et al., 2015). These VR-induced alterations in neural activity have been attributed to the loss of proprioceptive, vestibular, and motor efference signals, which are essential for generating coherent spatial representations (Stackman and Taube, 1997; Stackman and Herbert, 2002; Muir et al., 2009; Ravassard et al., 2013). These limitations have provided motivation for a shift toward hybrid designs that at least partially retain real-world locomotion while embedding dynamic sensory control by using computer-rendered visual cues that can be dynamically configured during constrained movement behavior (Jacobson et al., 2014; Kaupert et al., 2017; Stowers et al., 2017; Lester et al., 2020; Madhav et al., 2022). While these existing hybrid systems improve ethological validity, most are constrained to circular track locomotion (Stowers et al., 2017; Lester et al., 2020; Madhav et al., 2022), or strongly restrict linear motion (Kaupert et al., 2017), which greatly limits the repertoire of movement types sampled. Importantly, none of these hybrid approaches provide direct real-time programmatic control over available route motifs like that afforded by the Omniroute’s configurable gate system.

The Omniroute represents a largely open, real-world behavioral solution that provides the major benefits of a VR-based system and significantly extends the functionality of existing free-behavior apparatuses, enabling real-time coordination of routes, cues, and reward with fully programmable operation at sub-second scales. The findings presented demonstrate the system’s reliable closed-loop operation, centimeter-level reward targeting, and successful experimental application using a cue-based behavioral paradigm with changing spatial contexts. All of this is achieved while remaining compatible with electrophysiological investigations and normal animal welfare requirements. By providing VR-like control over sensory contingencies while preserving natural locomotion, the platform enables comparable real-world experiments that couple dynamic geometry with changing sensory inputs and flexible reinforcement in freely moving rodents. We anticipate adaptations to larger arenas and tasks requiring more dynamic event timing, and yet-to-be-conceived applications that exploit the full functionality of the system.

## Acknowledgements

This work could not have been completed without administrative support from Isaac Morgan, and animal care and training support from Rick Kornelsen. The following undergraduate researchers assisted A.W.L. with the assembly, testing and maintenance of the apparatus: Abhishek Dhir, Musa Habib, Nadira Djafri, Rebecca Alain, Matin Narimani and Gurnoor Kaur. Weilan Zhang assisted A.G.M. in rat training. This paper is dedicated to our fond memories of Rick Kornelsen.

## References

Aghajan ZM, Acharya L, Moore JJ, Cushman JD, Vuong C, Mehta MR (2015) Impaired spatial selectivity and intact phase precession in two-dimensional virtual reality. Nature Neuroscience 18:121–129.

Chen G, King JA, Burgess N, O’Keefe J (2013) How vision and movement combine in the hippocampal place code. Proceedings of the National Academy of Sciences of the United States of America 110:378–383.

Dombeck DA, Khabbaz AN, Collman F, Adelman TL, Tank DW (2007) Imaging large-scale neural activity with cellular resolution in awake, mobile mice. Neuron 56:43–57.

Holleman E, Mąka J, Schröder T, Battaglia F (2019) An incremental training method with automated, extendable maze for training spatial behavioral tasks in rodents. Scientific Reports 9:12589.

Jacobson TK, Ho JW, Kent BW, Yang F-C, Burwell RD (2014) Automated visual cognitive tasks for recording neural activity using a floor projection maze. Journal of Visualized Experiments e51316.

Jankowski MM, Polterovich A, Kazakov A, Niediek J, Nelken I (2023) An automated, low-latency environment for studying the neural basis of behavior in freely moving rats. BMC Biology 21:172.

Kaupert U, Thurley K, Frei K, Bagorda F, Schatz A, Tocker G, Rapoport S, Derdikman D, Winter Y (2017) Spatial cognition in a virtual reality home-cage extension for freely moving rodents. Journal of Neurophysiology 117:1736–1748.

Lester AW, Kapellusch AJ, Barnes CA (2020) A novel apparatus for assessing visual cue-based navigation in rodents. Journal of Neuroscience Methods 338:108667.

Lester AW, Kaur G, Djafri N, Madhav MS (2024) A modular gate system for autonomous control of rodent behavior. bioRxiv 2024.11.22.624912.

Lopes G, Bonacchi N, Frazão J, Neto JP, Atallah BV, Soares S, Moreira L, Matias S, Itskov PM, Correia PA, Medina RE, Calcaterra L, Dreosti E, Paton JJ, Kampff AR (2015) Bonsai: an event-based framework for processing and controlling data streams. Frontiers in Neuroinformatics 9:7.

Madhav MS, Jayakumar RP, Lashkari SG, Savelli F, Blair HT, Knierim JJ, Cowan NJ (2022) The Dome: a virtual reality apparatus for freely locomoting rodents. Journal of Neuroscience Methods 368:109336.

Mathis A, Mamidanna P, Cury KM, Abe T, Murthy VN, Mathis MW, Bethge M (2018) DeepLabCut: markerless pose estimation of user-defined body parts with deep learning. Nature Neuroscience 21:1281–1289.

Muir GM, Brown JE, Carey JP, Hirvonen TP, Della Santina CC, Minor LB, Taube JS (2009) Disruption of the head direction cell signal after occlusion of the semicircular canals in the freely moving chinchilla. The Journal of Neuroscience 29:14521–14533.

Muller RU, Kubie JL, Ranck JB (1987) Spatial firing patterns of hippocampal complex-spike cells in a fixed environment. The Journal of Neuroscience 7:1935–1950.

O’Keefe J, Dostrovsky J (1971) The hippocampus as a spatial map: preliminary evidence from unit activity in the freely moving rat. Brain Research 34:171–175.

O’Keefe J, Nadel L (1978) The hippocampus as a cognitive map. Oxford: Clarendon Press.

Olton DS, Samuelson RJ (1976) Remembrance of places passed: spatial memory in rats. Journal of Experimental Psychology: Animal Behavior Processes 2:97–116.

Packard MG, McGaugh JL (1996) Inactivation of hippocampus or caudate nucleus with lidocaine differentially affects expression of place and response learning. Neurobiology of Learning and Memory 65:65–72.

Pereira TD et al. (2022) SLEAP: a deep learning system for multi-animal pose tracking. Nature Methods 19:486– 495.

Porter BS, Olson JM, Leppla CA, Duvelle É, Bladon JH, van der Meer MAA, Jadhav SP (2025) Adapt-A-Maze: an open source adaptable and automated rodent behavior maze system. eNeuro ENEURO.0138–25.2025.

Quigley M, Conley K, Gerkey B, Faust J, Foote T, Leibs J, Wheeler R, Ng AY (2009) ROS: an open-source Robot Operating System In: Proceedings of the ICRA Workshop on Open Source Software, p5. Kobe, Japan: IEEE.

Ravassard P, Kees A, Willers B, Ho D, Aharoni D, Cushman J, Aghajan ZM, Mehta MR (2013) Multisensory control of hippocampal spatiotemporal selectivity. Science 340:1342–1346.

Sawatani F, Tamatsu Y, Ide K, Azechi H, Takahashi S (2022) Utilizing a reconfigurable maze system to enhance the reproducibility of spatial navigation tests in rodents. Journal of Visualized Experiments e64754.

Stackman RW, Herbert AM (2002) Rats with lesions of the vestibular system require a visual landmark for spatial navigation. Behavioural Brain Research 128:27–40.

Stackman RW, Taube JS (1997) Firing properties of head direction cells in the rat anterior thalamic nucleus: dependence on vestibular input. The Journal of Neuroscience 17:4349–4358.

Stowers JR, Hofbauer M, Bastien R, Griessner J, Higgins P, Farooqui S, Fischer RM, Nowikovsky K, Haubensak W, Couzin ID, Tessmar-Raible K, Straw AD (2017) Virtual reality for freely moving animals. Nature Methods 14:995–1002.

Thurley K, Ayaz A (2017) Virtual reality systems for rodents. Current Zoology 63:109–119.

White SR, Amarante LM, Kravitz AV, Laubach M (2019) The future is open: open-source tools for behavioral neuroscience research. eNeuro 6:ENEURO.0223–19.2019.

Wilson MA, McNaughton BL (1993) Dynamics of the hippocampal ensemble code for space. Science 261:1055– 1058.

Wood RA, Bauza M, Krupic J, Burton S, Delekate A, Chan D, O’Keefe J (2018) The honeycomb maze provides a novel test to study hippocampal-dependent spatial navigation. Nature 554:102–105.

